# An AI-Enabled Translational Drug Discovery Framework for NALCN Channelopathy: From High-Throughput Screening to Therapeutic Candidate

**DOI:** 10.64898/2026.10.08.757476

**Authors:** Nattaya Thongsepee, Jeremy A. Tanner, Srikanth Polusani, Alexandra Typou, Nicholas A. Clanton, Brandon B. Holmes, Philippe Lory, Donard S. Dwyer, Daohong Zhou, Arnaud Monteil, Yaxia Yuan

## Abstract

**Background:** More than 90% of rare diseases lack an approved therapy, partly reflecting economic barriers in developing treatments for small patient populations. A common bottleneck is costly and resource-intensive hit-to-lead prioritization. Congenital contractures of the limbs and face, hypotonia, and developmental delay (CLIFAHDD) is an ultra-rare neurodevelopmental disease often caused by *de novo* gain-of-function variants in the sodium leak channel NALCN, with no approved treatments.

**Methods:** We developed a cost and resource-efficient, AI-enabled translational drug discovery framework integrating phenotypic high-throughput screening (HTS), structural biology foundation models for hit-to-lead prioritization, and orthogonal experimental validation. We applied this framework for drug repurposing to identify NALCN inhibitors as therapeutic candidates for CLIFAHDD. We screened ∼1,200 approved drugs using a fluorescence-based membrane potential assay in doxycycline-inducible HEK293 cells stably expressing the NALCN channelosome. Boltz-2 modeling predicted binding affinity and mode to prioritize hits. Lead repurposing candidates were selected by a multidisciplinary expert review panel. Lead candidates were evaluated using patch-clamp electrophysiology in HEK293 cells expressing wild-type or CLIFAHDD-associated NALCN variants, primary mouse hippocampal neurons, and a *C. elegans* NALCN gain-of-function model.

**Results:** HTS identified 11 candidate NALCN inhibitors. Boltz-2 prioritized candidates based on predicted binding affinity and predicted binding within the NALCN pore, consistent with a direct pore-blocking mechanism. Aprepitant emerged as the lead candidate after multidisciplinary evaluation based on its predicted mechanism, pharmacologic properties, blood-brain barrier penetration, established safety profile, and regulatory approval in children and adults. Patch-clamp electrophysiology confirmed partial inhibition of NALCN-mediated sodium leak currents in HEK293 cells (IC₅₀=2.22 μM), with comparable potency against Ca_v_3.1 (IC_50_=1.63 µM). In primary mouse hippocampal neurons, aprepitant reduced sodium leak currents, hyperpolarized resting membrane potential, and decreased spontaneous firing. Aprepitant rescued hyperlocomotor and dystonic phenotypes in a *C. elegans* gain-of-function NALCN model. There was notable concordance between AI-predicted hits, phenotypic screening hits, and experimental validation.

**Conclusion:** These findings identify aprepitant as a lead therapeutic candidate for CLIFAHDD, and establish an AI-enabled translational drug discovery framework integrating phenotypic screening, structural modeling, and experimental validation for NALCN channelopathies. This scalable and resource-conscious strategy may accelerate hit-to-lead prioritization and drug discovery for rare and ultra-rare diseases lacking effective therapies.

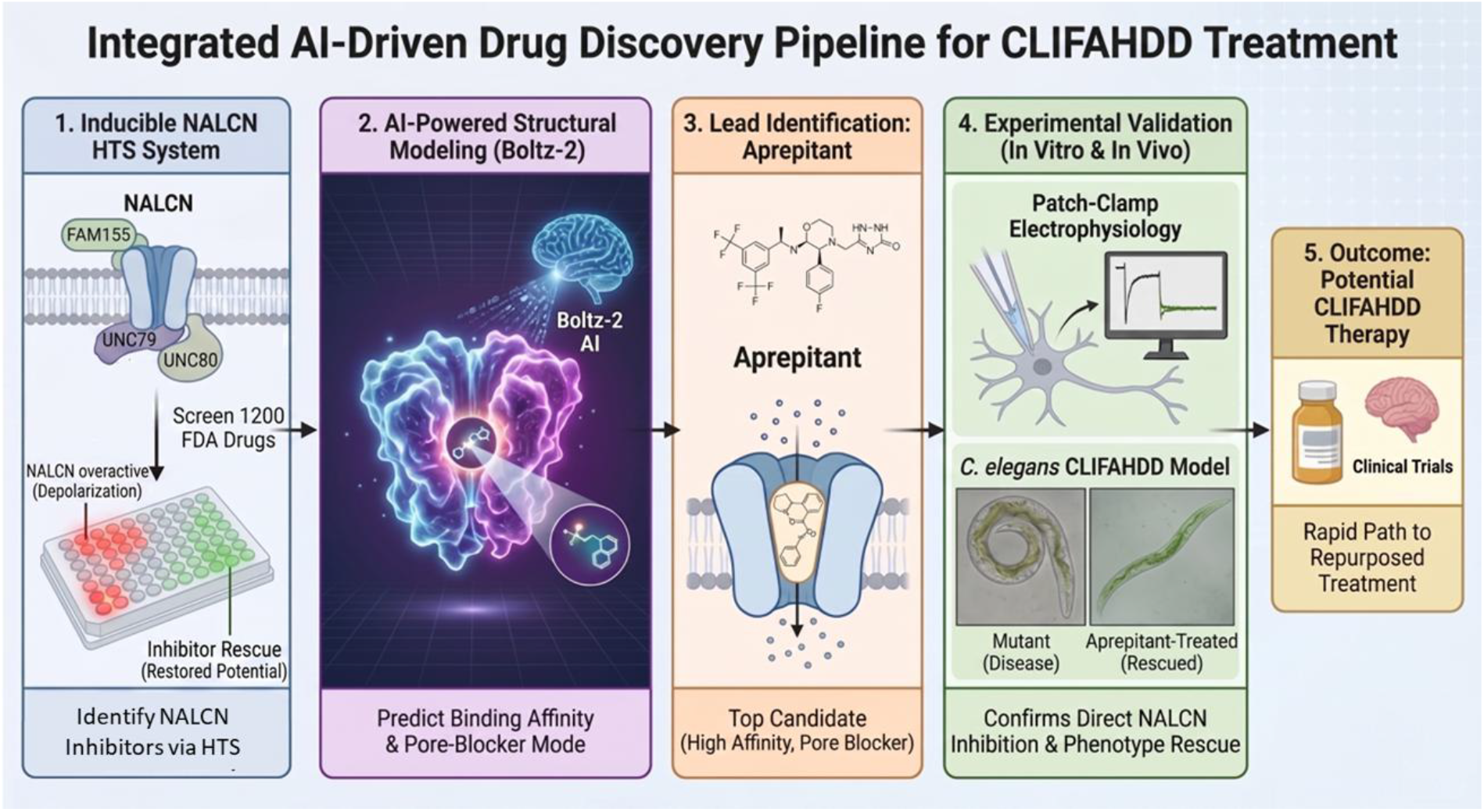

## Background

Rare diseases are increasingly recognized as a global public health priority[1]. Rare diseases collectively affect an estimated 3.5-5.9% of the global population and comprise more than 10,000 distinct diseases[2–4]. Most are pediatric, genetic, and ultra-rare[2–4], and the number of diagnosed patients continues to rapidly increase with expanding access to clinical genetic testing[1,4]. Despite this, approximately ∼95% of rare diseases lack an approved treatment option[5]. This therapeutic gap is driven in part by the high cost, lengthy timelines, and perceived cost-ineffectiveness associated with conventional drug discovery for diseases affecting very small patient populations, highlighting the need for more efficient therapeutic discovery strategies[1,6]. While drug repurposing offers a promising strategy for accelerating treatment development, identifying lead candidates for clinical trials remains challenging. Phenotypic drug repurposing screens often yield numerous hits and false positives as these compounds already possess diverse mechanisms of action, making hit-to-lead prioritization labor-intensive, costly, and a persistent bottleneck in translational drug discovery.

Congenital Contractures of the Limbs and Face, Hypotonia, and Developmental Delay (CLIFAHDD; OMIM #616266) is a severe, ultra-rare neurodevelopmental disease caused by heterozygous *de novo* missense mutations in *NALCN*[7–9]. Affected individuals present in early infancy with global neurodevelopmental delay, hypotonia, hyperkinetic movement disorders (*e.g.,* dystonia, ataxia), epilepsy, apnea, constipation, and sleep disturbances[7,10]. Individuals with CLIFAHDD experience substantial lifelong morbidity and an increased risk of early childhood mortality[7,10]. Despite its severe clinical burden, no approved treatments or clinical trials currently exist, reflecting a lack of prior drug discovery efforts and the broader therapeutic gap faced by most ultra-rare genetic diseases.

*NALCN* encodes the sodium leak channel (NALCN), a non-selective ion channel responsible for a background sodium (Na^+^) influx that plays a critical role in maintaining the resting membrane potential and regulating neuronal and cell excitability[11–15,9]. NALCN is the only member of a distinct branch of the four-domain ion channel family that includes the well-studied voltage-gated sodium and calcium channels[9,16]. The *NALCN* gene and subunits are highly conserved across species[17,18,14,9]. Accumulating evidence has demonstrated that NALCN and its ancillary subunits (UNC79, UNC80, FAM155A/B) contribute to fundamental physiological processes involving cell excitability, rhythmic processes, and pacemaking activity including neuronal firing[11], respiratory rhythm[11], circadian rhythm[19], nociception[20–22], gastrointestinal motility[23], hormonal secretion[24], and parturition[25],

In individuals with CLIFAHDD, pathogenic *de novo* variants can confer gain-of-function in NALCN characterized by pathological membrane depolarization and increased current density[26–28], Importantly, NALCN haploinsufficiency is well-tolerated, enabling a wide therapeutic window. While biallelic loss-of-function causes an ultra-rare recessive disease called IHPRF1, heterozygous carriers and individuals with monoallelic loss-of-function variants are asymptomatic[10,29–31]. As such, partial pharmacologic inhibition of NALCN represents an attractive therapeutic strategy for individuals with CLIFAHDD with gain-of-function variants. Consistent with this, pharmacologic inhibition of NALCN rescues disease phenotypes in *C. elegans* models of NALCN gain-of-function[32,33]. However, no clinically available NALCN inhibitors have been identified. A prior barrier to drug discovery and therapy development was the inability to generate robust, reproducible NALCN currents in recombinant expression systems suitable for high-throughput screening. We recently addressed this by developing an inducible HEK293 platform with stable expression of the complete NALCN channelosome, enabling reproducible functional measurements of NALCN current[33]. This created a new opportunity to systematically screen therapeutic candidates for CLIFAHDD.

Here we developed an AI-enabled drug discovery framework that integrates phenotypic high-throughput screening (HTS) with structure-based computational hit-to-lead prioritization to identify a lead repurposed drug candidate for CLIFAHDD, which we then validated using *in vitro* and *in vivo* NALCN models. An international, multidisciplinary drug discovery consortium was facilitated by the non-profit and patient advocacy organization, Channeling Hope Foundation, to design a cost and resource-efficient drug discovery strategy for CLIFAHDD. For hit discovery, we used the inducible HEK293 platform expressing the NALCN channelosome to develop a membrane depolarization assay, then screened a drug-repurposing library of ∼1200 compound enriched for FDA-approved drugs to identify hits. We then applied the recent structural biology foundation model Boltz-2[34] to prioritize compounds based on the predicted binding affinity and binding mode to directly inhibit the NALCN central pore. Separately and retrospectively, we employed Boltz-2 to screen the same library of ∼1200 repurposed drugs in order to benchmark the model against the phenotypic result; the predicted hits and the lead repurposing candidate were concordant with the top hits from phenotypic screening. The lead candidate that emerged was aprepitant, a neurokinin-1 (NK-1) receptor antagonist with blood-brain barrier penetrance and FDA-approval for use in children and adults with well-established safety and pharmacokinetic profile. Next, we validated aprepitant and evaluated other NK-1 receptor inhibitors using *in vitro* models with patch-clamp electrophysiology to confirm partial NALCN current inhibition in HEK293 and hippocampal neurons, and then *in vivo* with a *C. elegans* gain-of-function model to confirm phenotype rescue. These supported aprepitant as a promising lead therapeutic candidate, now advancing to a phase I clinical trial for CLIFAHDD. Overall, this demonstrated a scalable drug discovery approach leveraging AI to enable more cost-efficient hit-to-lead prioritization that can be applied to accelerate drug discovery and repurposing for channelopathies and other rare diseases without treatments.

## Methods

### Overview

We developed and validated an integrated, AI-enabled drug discovery pipeline to identify therapeutic candidates for CLIFAHDD caused by gain-of-function variants in NALCN. The pipeline combined (a) a cell-based phenotypic assay for HTS using an inducible HEK293 cellular platform expressing the recombinant NALCN channelosome to identify hits; (b) AI-based structural modeling to prioritize hits based on predicted mechanism and binding with NALCN to select leads; (c) lead validation *in vitro* using patch-clamp electrophysiology in recombinant cells and primary hippocampal neurons; (d) lead validation *in vivo* with a *C. elegans* disease model. HEK293 cells were selected for primary screening because they are amenable to genetic engineering, scalable for HTS, and lack contamination from other ion channels. During assay development, the membrane potential assay was initially designed and piloted using transient transfection of plasmids encoding the complete NALCN channelosome (NALCN, UNC79, UNC80, and FAM155A). Once an inducible HEK293 cell line with stable expression of the NALCN channelosome became available, all HTS experiments were performed using the stable cell line.

### Cell Lines

The generation and characterization of inducible HEK293 cell lines stably expressing wild-type NALCN with its three subunits UNC79, UNC80 and FAM155A were recently described[33]. Briefly, HEK293T (tsA201) cells were obtained from the European Collection of Authenticated Cell Cultures (ECACC #96121229 respectively). Cell identity was confirmed by Short Tandem Profiling (STR) profiling. The cells were routinely tested for the absence of any mycoplasma contamination. For HEK293 cells, the following antibiotics were added in the culture media: Hygromycin (200 μg/ml), Puromycin (1 μg/ml), Blasticidin (10 μg/ml) and Geneticin (400 μg/ml; G418 sulfate). NALCN expression was later induced by adding Doxycycline Hyclate ReadyMade™ (Dox).

### High Throughput Screening for Hit Discovery

A fluorescence-based membrane potential assay was developed and optimized for HTS on 384-well plates using the inducible HEK293 NALCN channelosome line. Membrane depolarization was measured using a live-cell membrane potential dye (Molecular Devices) with fluorescence recorded on the Fluorometric Imaging Plate Reader (FLIPR) Penta HTS System. To minimize potential interference from high Dox concentrations on the cellular system, we evaluated the effects of induction with 0.1, 0.5, 1, and 2 μg/mL Dox on membrane depolarization, ultimately selecting 1 µg/mL as an optimal concentration for screening (**Fig. S1**). HEK293 cells were seeded onto poly-L-lysine-coated 384-well plates at 5000 cells per well in DMEM supplemented with GlutaMAX (Gibco, 10566-016) and Dox (1 μg/ml). After 72 hours, each of the ∼1,200 drug-repurposing compounds from Prestwick Chemical Library enriched for FDA-approved drugs were tested at a final concentration of 10 μM in 0.02% DMSO with 30 minutes of compound exposure and 45 minutes of membrane potential dye exposure. The positive control was 10 mM CaCl_2_, a non-specific and direct NALCN channel inhibitor[18,26]. The negative controls were DMSO-treated induced cells and non-induced cells. Compounds that reduce membrane potential depolarization intensity by 60% or more are presented as hits. Primary screening results obtained with the FLIPR system were independently confirmed by high-content fluorescence imaging using the Opera Phenix platform.

### AI-Based Protein-Ligand Modeling and Binding Affinity Prediction for Hit-to-Lead Prioritization

Boltz-2, an AI-based protein-ligand structure prediction framework[34], was used to predict the binding affinities and binding modes of all HTS hits. Boltz-2 was selected for predicting protein-ligand binding based on prior evidence of achieving high accuracy in both binding affinity prediction and structural modeling. Its co-folding approach predicts protein structure and ligand pose simultaneously, making it particularly well-suited for proteins exhibiting conformational change effects, such as NALCN. The prediction was performed under default parameters, utilizing the full-length sequence of human NALCN and small molecules converted into SMILES format. For downstream data visualization, the predicted binding affinities were converted into LogIC_50_ units. The binding pose with the highest predicted affinity was selected as the representative model and visualized using PyMOL (Schrödinger). To further evaluate the performance of Boltz-2 relative to experimental validation, the same methodology was later applied to predict binding for (a) the overall library of ∼1,200 FDA-approved drugs used in phenotypic screening to NALCN; (b) additional NK-1 receptor antagonists to NALCN, and (c) lead candidate to Ca_v_3.1 and Na_v_1.5 channels to evaluate predicted selectivity for NALCN among four-domain ion channels.

### Multidisciplinary Lead Repurposing Candidate Selection

A multidisciplinary expert review panel was convened to review and prioritize hits as lead repurposing candidates for further validation. Hit compounds were evaluated based on evidence from phenotypic HTS, AI-based structural modeling, published NALCN and disease biology, blood-brain barrier permeability, pharmacokinetics and pharmacodynamic properties, safety profiles, route of administration, clinical evidence and experience, FDA-approved indications and labeling, and feasibility for clinical translation. Lead candidates for further validation were selected based on the totality of this evidence.

### In Vitro Validation with Patch Clamp Recordings in HEK293 Cells and Mouse Primary Hippocampal Neurons

The lead candidate and mechanistically related compounds (NK-1 receptor antagonists) were advanced to orthogonal functional validation by whole-cell patch clamp electrophysiology. *In vitro* validation was designed to assess (a) NALCN current inhibition in HEK293 cells stably expressing the wild-type NALCN channelosome; (b) generalizability by evaluating NALCN current inhibition in HEK293 cells stably expressing CLIFAHDD patient-derived gain-of-function variants; (c) selectivity by evaluating compound effects on Ca_v_3.1 and Na_v_1.5 currents in HEK293 cells; (d) inhibition of endogenous NALCN current in mouse primary hippocampal neurons. NK-1 receptor antagonists evaluated included aprepitant (Sigma-Aldrich SML2215), rolapitant (Selleckchem Z815946), netupitant (MedChemExpress HY-16346) and tradipitant (MedChemExpress HY-16732). Additional details are provided in the Supplementary Methods.

#### NALCN Current Inhibition Assays

HEK293 cells with stable expression of wild-type NALCN channelosome were prepared as described above. NALCN expression was induced by adding Dox at a concentration of 0.1 μg/ml in the culture media and cells were locked in the M phase of the cell cycle with nocodazole (100 ng/ml). Cells were dissociated with Versene prior to recording. Whole-cell patch clamp recordings of macroscopic sodium leak (NALCN) currents were performed at room temperature using an Axopatch 200B amplifier (Molecular Devices). Data were acquired using the pCLAMP10 software (Molecular Devices) and an Axon Digidata 1440A digitizer (Molecular Devices) at 10 kHz. Patch pipettes were pulled from 1.5/086 borosilicate glass capillaries. Intracellular and extracellular recording solutions are described in the Supplementary Methods.

#### Generalizability Assays

The lead candidate was tested for NALCN current inhibition in two HEK293 cell-lines stably expressing CLIFAHDD patient-derived NALCN gain-of-function variants p.Y578S and p.D600H. These pathogenic variants have previously been shown to have increased NALCN current density in recombinant expression systems, and altered sensitivity and degrees of blockade to pharmacological compounds compared to the wild-type channel[26,28,33]. Methods for generation, characterization, and preparation of these stable cell lines have been recently detailed[33]. Electrophysiological recordings were performed using the same methods described above for wild-type NALCN.

#### Selectivity Assays

The lead candidate and an analog compound were tested for off-target inhibition of two four-domain voltage-gated ions channels, Na_v_1.5 and Ca_v_3.1, in HEK293T cells. For transient functional expression of Na_v_1.5 and Ca_v_3.1 channels, 1.9 µg of plasmid DNA supplemented with 0.1 µg of pcDNA3-GFP was used to transfect HEK293T cells in 35 mm Petri dishes using the jetPEI reagent according to the manufacturer’s protocol. Whole-cell patch clamp recordings were performed using the same electrophysiological configuration described for NALCN current recordings. Channel-specific intracellular and extracellular recording solutions are detailed in the Supplementary Methods.

#### Mouse Primary Hippocampal Neurons

Primary hippocampal neurons were prepared from newborn C57BL/6J pups as previously described[35]. Cells were cultured under standard conditions and maintained until DIV14–20 for electrophysiological recordings. Hippocampal cultures from >3 independent litters were used for whole-cell patch clamp recordings between DIV14 and DIV20, after synapse maturation. Additional details on neuronal culture and electrophysiology are available in the Supplementary Methods.

### In Vivo Validation in C. elegans with Gain-of-Function Mutation in NALCN

For *in vivo* orthogonal validation, we used a *C. elegans* mutant, *unc-77(e625)*, which harbors a gain-of-function mutation (p.A717V located in the sixth transmembrane segment of the second domain; distinct from patient-derived variants modeled in HEK293 cells above) in the NALCN ortholog, *nca-1*[32]. NALCN and NCA-1 share a high degree of protein sequence homology, particularly within the transmembrane and pore-forming domains[17]. These variants display a reproducible gain-of-function phenotype characterized by hyperactive locomotion, exaggerated body bending, and excessive coiling. This phenotype has been previously validated as a quantitative pharmacodynamic readout of NALCN inhibition[32].

#### Drug Concentrations

Compound concentrations were selected based on prior *C. elegans* studies with NK-1 receptor antagonists by our group[33] and with other psychotropic drugs[36,37]. Consistent with prior reports, micromolar levels (20-160 mM) of drug were used on agar plates to account for limited drug uptake through the nematode cuticle and restricted tissue distribution to achieve pharmacologically relevant *in vivo* target exposure[38]. Additional information is provided in the Supplementary Methods.

#### Foraging Assay

The hyperactive foraging phenotype was quantified using a previously described and validated behavioral assay[32]. Briefly, animals were incubated on control (dilute acetic acid plus dimethyl sulfoxide, DMSO) or drug-containing agar plates for 2 hours before transfer to bacteria-free assay plates. Foraging behavior was quantified over a 3-minute observation period by counting the total number of sharp “omega” turns (head touching body), pirouettes, and reversals. Each experiment was repeated at least twice, and data was obtained from a minimum of 10 animals per condition. Detailed plate preparation and drug administration procedures are described in the Supplementary Methods.

#### Coiling Assay

Coiling behavior was quantified using a previously described assay[32]. Control (DMSO) and drug plates were prepared as described above and 10-15 animals were transferred to each plate. After 2 hours, we examined whether animals displayed the S-posture typical of wild-type animals or an atypical coiling phenotype with the head touching the body. The number of animals in a coil or omega posture was counted along with the total number of animals on the plate to determine the percent coiling. Each experiment was replicated at least 3 times for analyses.

### Data Analyses and Availability

Electrophysiological data were analyzed using pCLAMP10 (Molecular Devices), Statistical analyses were performed using GraphPad Prism 10 and Microsoft Excel. Data are presented as the mean ± SEM unless otherwise indicated, with *n* indicating the number of cells. Normality was assessed using the Shapiro-Wilk test. Statistical tests used in analyses are specified in the corresponding figure legends.

## Results

### Membrane Potential Assay Developed for High-Throughput Screening

Because the NALCN channel generates a constitutive sodium leak current that depolarizes the resting membrane potential[9,11,15,18,24,26], we developed a scalable fluorescence-based membrane potential assay using an inducible HEK293 cell line stably expressing the NALCN channelosome. Induction of channelosome expression with doxycycline produced a quantifiable and reproducible increase in fluorescence, consistent with membrane depolarization, compared with uninduced controls on the FLIPR system (**Fig. S1**). NALCN inhibition with the positive control CaCl₂, a non-selective inhibitor of NALCN current, significantly reduced the fluorescence signal, consistent with membrane repolarization, whereas the negative control (DMSO) had no effect (**Fig. 1A**). Fluorescence changes measured by FLIPR were confirmed by high-content imaging (data not shown). Together, these results established a reproducible, induction-dependent assay with sufficient dynamic range to identify inhibitors of NALCN-mediated membrane depolarization at a single screening concentration.

**Fig. 1:**
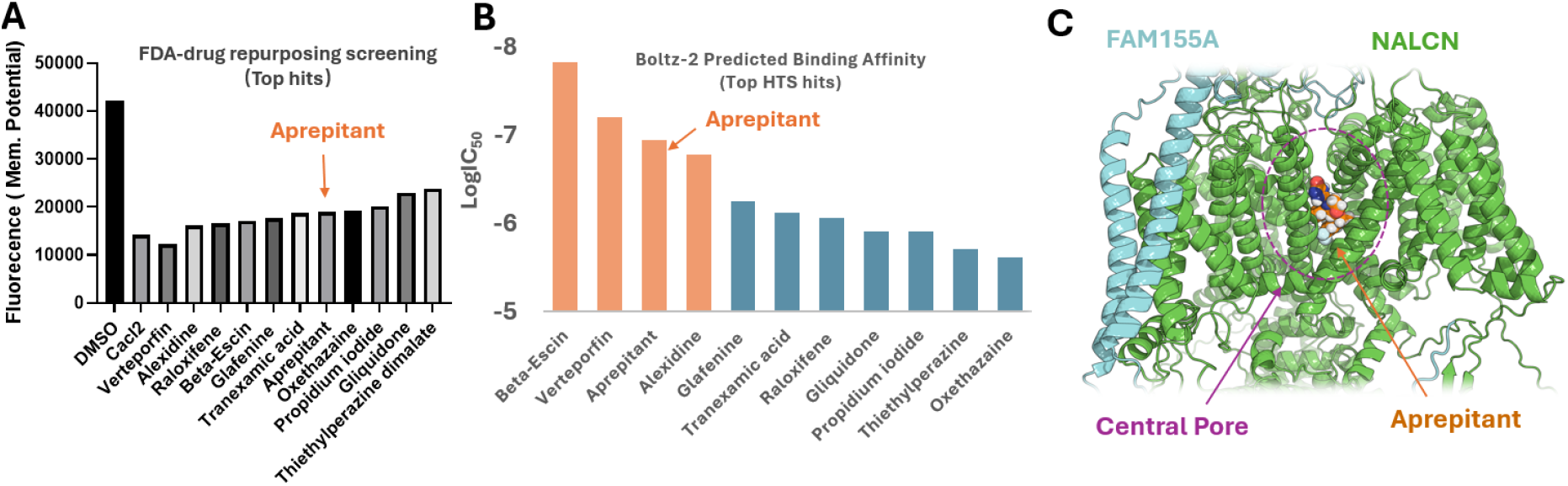
Implementation of a high-throughput screening assay to identify NALCN modulators. **(A)** Results of phenotypic high-throughput screening on ∼1,200 FDA drugs, showing the top 11 hits with the negative (DMSO) and positive (CaCl_2_) control. **(B)** Use of Boltz-2 to predict the binding strength of the top 11 phenotypic hits to NALCN. A LogIC_50_ value of -6 represents an IC_50_ = 1 μM; -7 represents an IC_50_ = 100 nM. **(C)** The predicted binding mode of aprepitant to NALCN using Boltz-2 is within the central ion-conducting pore in a direct pore-blocking mechanism

### Phenotypic High-Throughput Screening Identifies Candidate NALCN Inhibitors

We screened the Prestwick Chemical Library, a drug-repurposing library enriched for FDA-approved and biologically active substances (∼1,200 compounds), using the fluorescence-based membrane potential assay to identify inhibitors of NALCN-mediated membrane depolarization. Compounds were ranked by their ability to reduce the fluorescence signal, indicative of membrane repolarization. Eleven compounds significantly decreased fluorescence by at least 60% compared with the DMSO control (p < 0.05) and were prioritized as primary hits for further validation (**Fig. 1A**).

### AI-Based Binding Affinity and Binding Mode Prediction for Hit-to-Lead Prioritization

To prioritize the 11 primary screening hits, we employed Boltz-2 to predict ligand binding affinities and binding modes for NALCN. The four compounds with the highest predicted binding affinities were beta-escin, verteporfin, aprepitant, and alexidine (**Fig. 1B**). To further prioritize hits, we sought compounds predicted to directly bind in the NALCN central pore as one promising mechanism to identify inhibitors. For this, we used Boltz-2 to model the binding poses of NALCN-compound complexes, allowing us to identify compounds predicted to bind within the NALCN central pore as direct pore blockers. Aprepitant and alexidine were predicted to occupy and partially occlude the central, ion-conducting pore (**Fig. 1C**). While beta-escin and verteporfin were predicted to bind with high affinity, their bulky scaffolds were predicted to sterically preclude central pore entry.

To evaluate concordance between AI-based predictions and the phenotypic screen, we retrospectively applied Boltz-2 to the entire Prestwick Chemical Library (∼1,200 compounds). Four of the 11 primary hits on HTS ranked within the top 30 compounds for predicted NALCN binding affinity by Boltz-2. These included the two compounds that produced the greatest fluorescence suppression on HTS (verteporfin and alexidine), and aprepitant, which was ultimately advanced as the lead candidate (**Fig. S2**). Among these 30 compounds, nine were predicted to bind within the NALCN central pore, including aprepitant.

### Multidisciplinary Prioritization of Lead Candidates

The 11 primary hits were then evaluated by a multidisciplinary review panel integrating phenotypic screening results, AI-predicted binding affinity and binding mode, relevance to NALCN biology, blood-brain barrier permeability, pharmacokinetics, clinical safety data, regulatory approval status, and translational feasibility. Based on these criteria, aprepitant, an FDA-approved NK-1 receptor antagonist, emerged as the lead repurposing candidate. Converging evidence supporting the translational potential of aprepitant included: (a) AI-predicted high-affinity binding in a direct, partial pore-blocking mechanism; (b) reproducible inhibition in orthogonal phenotypic assays (*data not shown*); (c) established blood-brain barrier permeability; (d) FDA-approval for use in pediatric (age >6 months) and adult populations; (e) well-characterized pharmacokinetic and safety profiles with a wide therapeutic index; (f) and independent evidence that some NK-1 receptor antagonists inhibit NALCN[22,23,39–52]. Consistent with a direct effect on NALCN, wild-type HEK293 cells do not endogenously express NK-1 receptors[53], indicating the observed NALCN inhibition is independent of NK-1 receptor signaling.

Given prior evidence implicating NK-1 receptor antagonists as NALCN inhibitors, three additional FDA-approved compounds from this class—rolapitant, netupitant, and tradipitant— were selected alongside aprepitant for further pharmacological validation.

### Initial Evaluation of Lead Candidates

To verify that aprepitant alters membrane potential through NALCN, we tested its effect in the inducible-HEK293 cells in the presence and absence of Dox to induce NALCN current using the fluorescence-based membrane potential assay (**Fig. 2**). In uninduced cells, aprepitant had no effect on membrane potential. After Dox induction, aprepitant attenuated NALCN-mediated depolarization, supporting that its activity in this assay is dependent on NALCN expression.

**Fig. 2:**
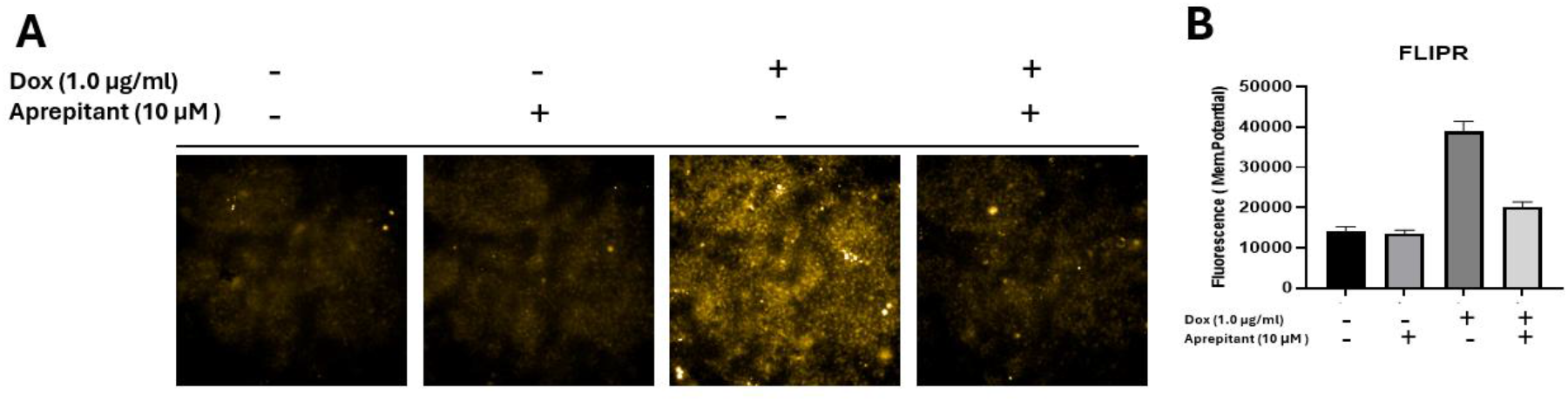
Effect of aprepitant with or without Dox treatment. **(A)** Comparison of cellular responses to aprepitant in the presence or absence of Dox induction. Lower fluorescence indicates a lower resting membrane potential. Higher fluorescence signifies a higher resting membrane potential (membrane depolarization) indicating NALCN channelosome expression. **(B)** Quantification of cellular fluorescence intensity. In the absence of Dox induction, aprepitant treatment exhibits no effect on the resting membrane potential. However, upon Dox induction, aprepitant significantly attenuates the membrane depolarization induced by NALCN upregulation.

We next used Boltz-2 to predict the binding affinities and modes of NK-1 receptor antagonists (**Fig. 3; Fig. S3**). Aprepitant had the highest predicted binding affinity among the NK-1 receptor antagonists tested (**Fig. 3A**), suggesting that aprepitant possesses specific structural features favorable for NALCN engagement. Rolapitant, the compound most structurally similar to aprepitant, showed overlap of the shared 3,5-bis(trifluoromethyl)phenyl moieties within the predicted NALCN binding pose (**Fig. 3B, C**). *In silico* structure-activity relationship (SAR) analysis suggested that aprepitant’s specific morpholine-acetal scaffold may be particularly favorable for NALCN binding. Both compounds were predicted to occupy the central ion-conducting pore of NALCN, consistent with a direct pore-blocking mechanism, though significant deviations were observed in the orientations of the remaining structural regions.

**Fig 3.**
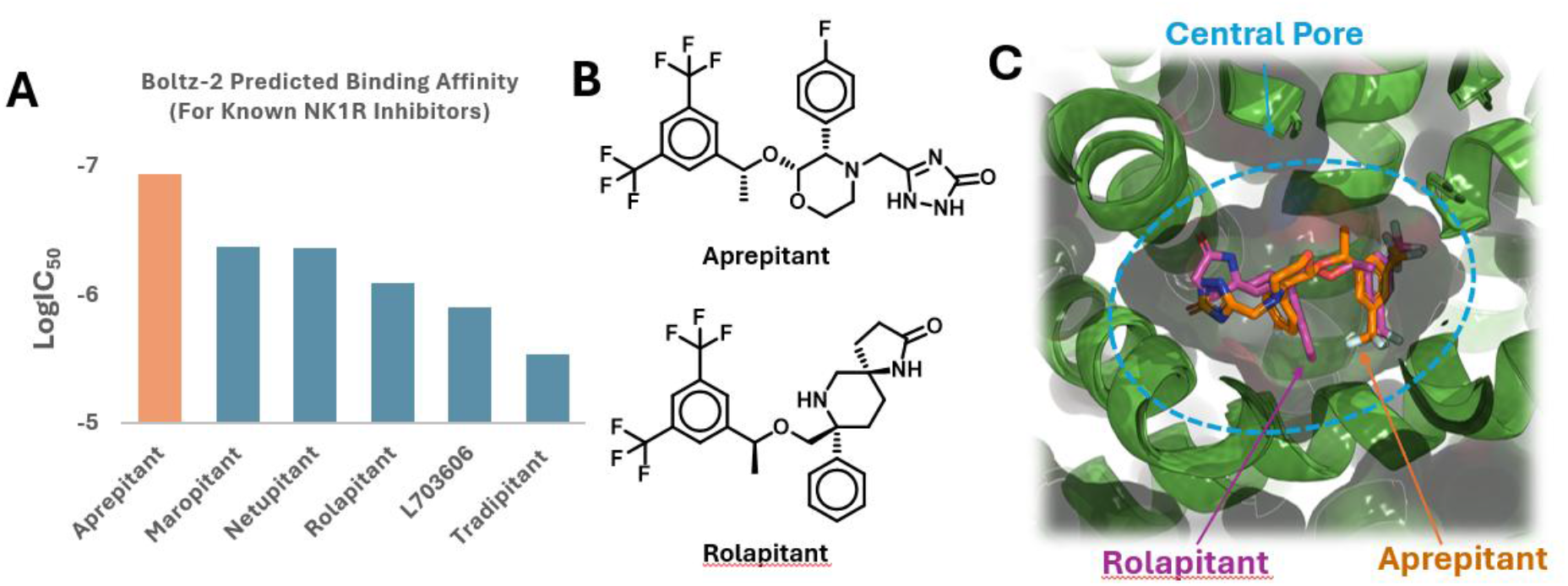
Binding modes of Aprepitant and other NK1R inhibitors. **(A)** Boltz-2 predicted binding affinities of various NK-1R inhibitors, indicating that aprepitant exhibits the highest binding strength. **(B)** Chemical structures of aprepitant and its most structurally similar marketed counterpart, rolapitant. **(C)** Structural superimposition of the predicted binding modes of aprepitant and rolapitant within NALCN. The central pore is delineated by blue dashed lines.

To assess predicted ion-channel selectivity, we employed Boltz-2 to model aprepitant binding to NALCN and to related four-domain channels Ca_v_3.1 and Na_v_1.5 (**Fig. S4**). Aprepitant was predicted to bind within the central pores of all three channels, but with distinct channel-specific poses. Specifically, the predicted conformation of aprepitant within Ca_v_3.1 was relatively similar to NALCN, whereas its conformation within Na_v_1.5 differed substantially. Consistent with these structural predictions, Boltz-2 predicted comparable binding affinities for NALCN and Ca_v_3.1, but lower affinity for Na_v_1.5.

### In Vitro Validation of NALCN Inhibition in HEK293 cells

We next evaluated aprepitant and other NK-1 receptor antagonists using whole-cell patch clamp recordings in HEK293 cells stably expressing the wild-type NALCN channelosome. At 20 µM, aprepitant and rolapitant significantly and partially inhibited NALCN currents under low extracellular Ca²⁺ conditions at a membrane potential of −40 mV (**Fig. 4**). Similar inhibition was observed under 2 mM extracellular Ca²⁺ conditions (**Fig. S5**). Dose–response curves were subsequently generated at −40 mV for aprepitant and rolapitant under 2 mM extracellular Ca²⁺ conditions (**Fig. 5**). Aprepitant inhibited NALCN current with an IC₅₀ of 2.22 µM and maximal inhibition of 36%, whereas rolapitant exhibited similar potency (IC₅₀ = 3.44 µM) and greater maximal inhibition (66%). Netupitant and tradipitant exhibited substantially lower potency (IC₅₀ = 14.7 and 16.9µM, respectively) and produced partial maximal inhibition (41% and 44%, respectively) (**Fig. S6**). These electrophysiological findings validated the primary phenotypic screening results for aprepitant and were consistent with AI-predicted binding affinities and partial occlusion of the central pore used for hit prioritization among HTS hits and among NK-1 receptor antagonists. Together, these results support aprepitant and rolapitant as partial inhibitors of NALCN.

**Fig. 4:**
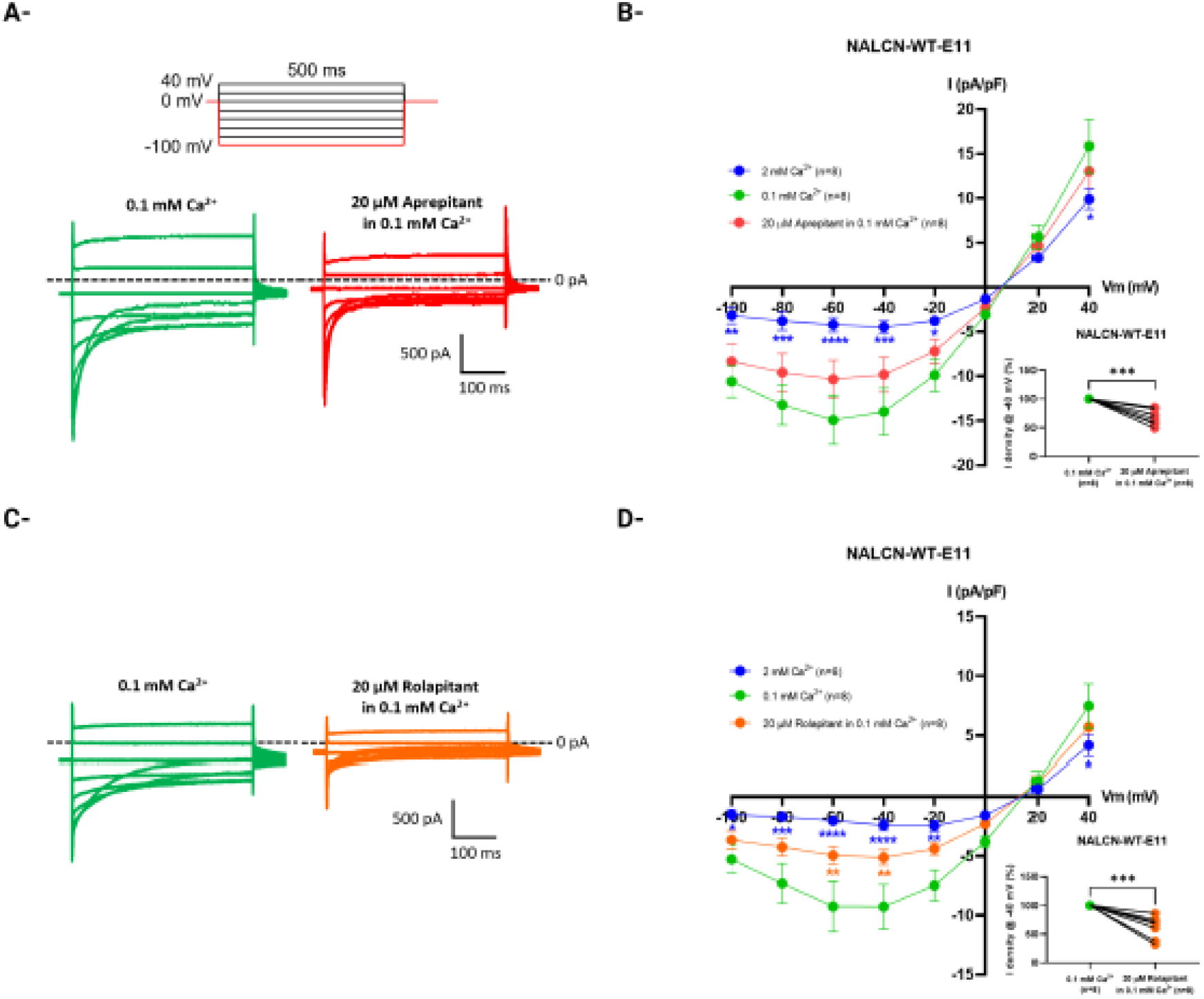
Modulation of the NALCN-mediated Na⁺ leak current by aprepitant and rolapitant. The effects of aprepitant and rolapitant were assessed at 20 µM on the NALCN current elicited by a voltage-step protocol in the monoclonal HEK293 cell line stably expressing the wild-type NALCN channelosome [33]. **A)** and **C)** Representative current traces recorded in 0.1 mM extracellular Ca²⁺ before and after application of aprepitant and rolapitant, respectively. **B)** and **D)** Current density–voltage (I/V) relationships derived from **A)** and **C)**, respectively. Insets show paired analyses of current density at −40 mV. Statistical analysis: two-way ANOVA with Dunnett’s multiple comparisons test (I/V relationships) and paired t-test (insets). ****p < 0.0001, ***p < 0.001, **p < 0.01, *p < 0.05.

**Fig. 5:**
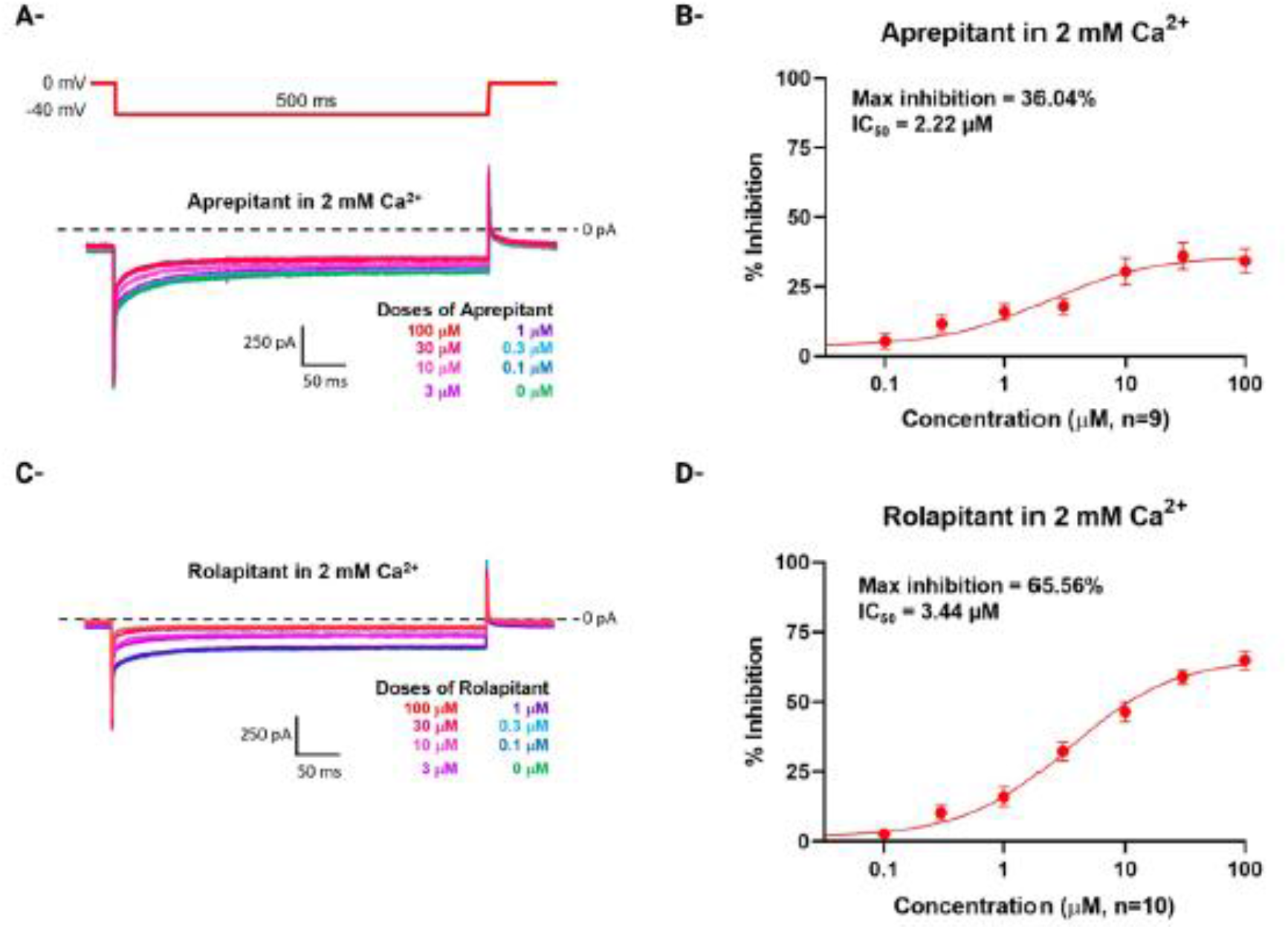
Concentration–response characterization of NALCN current inhibition by aprepitant and rolapitant in 2 mM extracellular Ca²⁺. The inhibitory properties of aprepitant and rolapitant on the NALCN current (apparent affinity and maximal inhibition) were characterized using a voltage step to −40 mV from a holding potential of 0 mV. **A)** and **C)** Representative current traces showing NALCN current inhibition following application of increasing concentrations (0.1–100 µM) of aprepitant and rolapitant, respectively. **B)** and **D)** Concentration–response relationships for inhibition of the NALCN current by aprepitant and rolapitant, respectively.

Notably, repeating dose–response assays under low extracellular Ca²⁺ (0.1mM) reduced the apparent potency of both aprepitant (IC_50_ = 4.62µM) and rolapitant (IC_50_ = 11.9µM) relative to standard Ca²⁺ conditions (2mM), while maximal inhibition was essentially unchanged for either compound (49% and 69%, respectively; **Fig. S7**). This selective effect on potency, with efficacy preserved, indicates that extracellular Ca²⁺ — itself a pore blocker of NALCN — may facilitate, rather than compete, with the binding of these compounds. A simple competitive mechanism, in which Ca²⁺ and the compounds occupy mutually exclusive sites, would predict the opposite result (increased potency at low Ca²⁺). We instead hypothesize that Ca²⁺ occupancy at or near the pore is required to stabilize a high-affinity conformation of the compounds’ binding site, either directly by contributing to the binding determinants, or indirectly by favoring a channel gating state that these compounds bind more readily. Both compounds remain partial inhibitors of NALCN current across the extracellular Ca²⁺ range tested.

### Aprepitant Generalizability for NALCN Inhibition in Pathogenic De Novo Variants

Because pathogenic NALCN gain-of-function variants associated with CLIFAHDD can exhibit altered pharmacological responses relative to the wild-type channel[33], we evaluated the effects of aprepitant on two *de novo* gain-of-function variants, p.Y578S and p.D600H, in HEK293 cells stably expressing the human NALCN channelosome (**Fig. 6**). Aprepitant inhibited both variant channels with apparent potencies comparable to wild-type NALCN (wild-type: IC₅₀ = 2.22 µM; p.Y578S: 1.39 µM; p.D600H: 2.33 µM) (**Fig. 6B, D**). The maximal inhibition of aprepitant at −40 mV was reduced for both variants (∼14%) relative to wild-type NALCN (∼36%). These results suggest that pathogenic variants of NALCN may exhibit altered pharmacological sensitivity relative to the wild-type channel. Importantly, because CLIFAHDD results from heterozygous *de novo* gain-of-function variants, patients express both wild-type and mutant NALCN channels. Thus, partial inhibition of both the wild-type channel and the mutant channel are expected to contribute to overall reduction of NALCN-related hyperexcitability in individuals with gain-of-function variants.

**Fig. 6:**
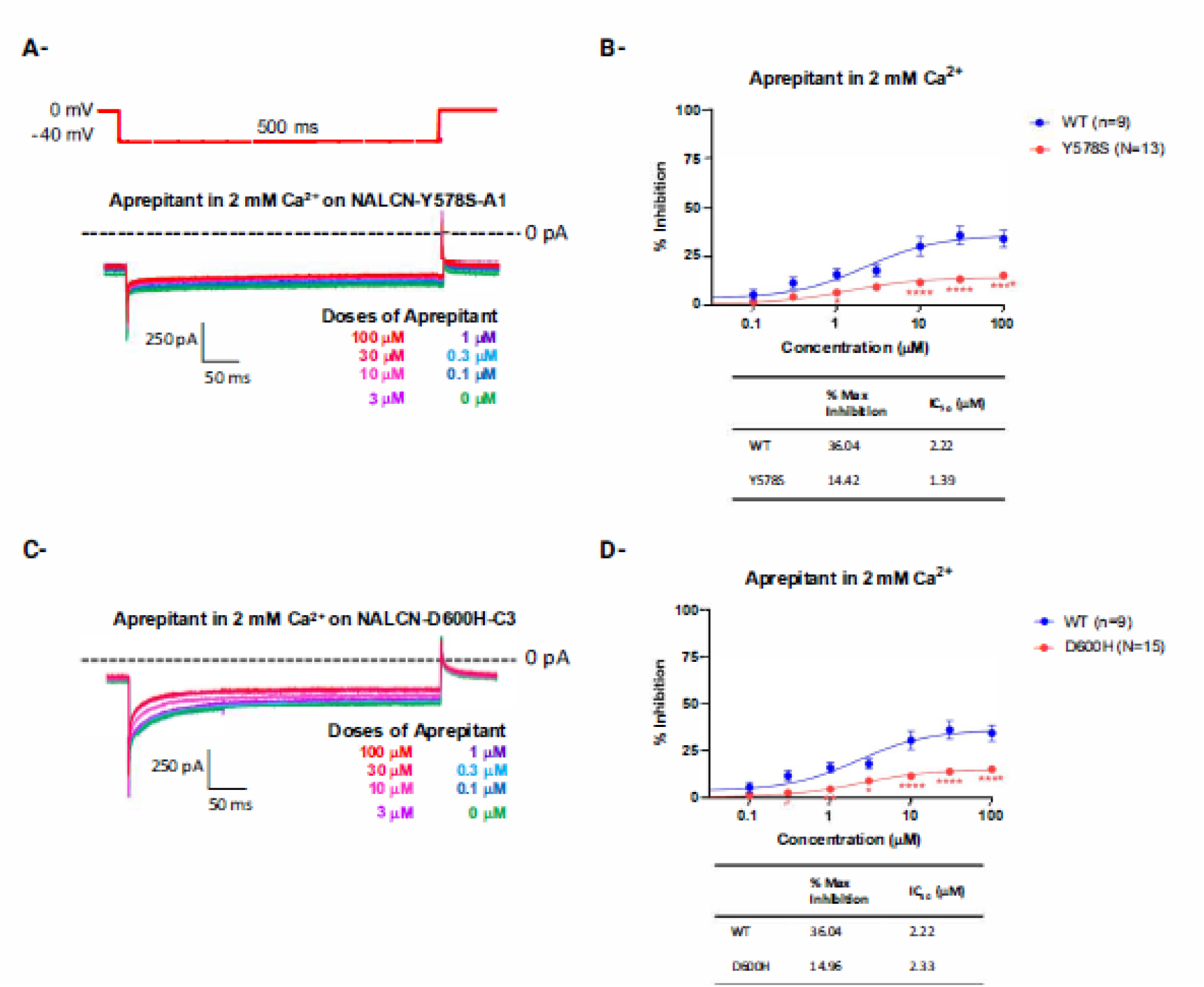
Concentration–response characterization of NALCN current inhibition by aprepitant in two gain-of-function variants. The inhibitory properties of aprepitant (apparent affinity and maximal inhibition) were assessed in two monoclonal HEK293 cell lines expressing gain-of-function *de novo* pathogenic variants of NALCN, using a voltage step to −40 mV from a holding potential of 0 mV. **A)** and **C)** Representative current traces for the p.Y578S and p.D600H variants, respectively, showing current inhibition following application of increasing concentrations of aprepitant (0.1–100 µM). **B)** and **D)** Concentration–response relationships for inhibition of NALCN currents by aprepitant for the p.Y578S and p.D600H variants, respectively.

### Selectivity Assays for Inhibition of Ca_v_3.1 and Na_v_1.5 Currents

To assess the selectivity of aprepitant and rolapitant for NALCN, we examined their effects on two voltage-gated ion channels sharing the four-domain topology: the T-type calcium channel Ca_v_3.1 (**Fig. 7**) and the cardiac sodium channel Na_v_1.5 (**Fig. 8**). Both compounds inhibited Ca_v_3.1 currents with greater apparent potency than NALCN in a voltage-independent manner (aprepitant: IC_50_ = 1.63 µM, max inhibition = 93%; rolapitant: IC_50_ = 0.02µM, max inhibition = 66%) (**Fig. 7**). Against Na_v_1.5, both compounds produced near-complete inhibition (83– 99%), but exhibited markedly different potencies. Aprepitant inhibited Na_v_1.5 with an IC₅₀ of ∼40.5 µM, approximately 18-fold lower than its potency against NALCN, whereas rolapitant was substantially more potent with an IC₅₀ of ∼0.057 µM at −20 mV (**Fig. 8**). Collectively, these results indicate that aprepitant and rolapitant exhibit distinct selectivity profiles across related four-domain ion channels, rather than selective inhibition of NALCN. The substantially lower potency of aprepitant against Na_v_1.5 suggests that pharmacologically relevant concentration sufficient to inhibit NALCN are unlikely to produce significant Na_v_1.5 block. These electrophysiological findings were consistent with AI-based structural modeling predictions that aprepitant binds within the central pore of all three channels, with comparable predicted binding affinity for NALCN and Ca_v_3.1, but lower affinity for Na_v_1.5.

**Fig. 7:**
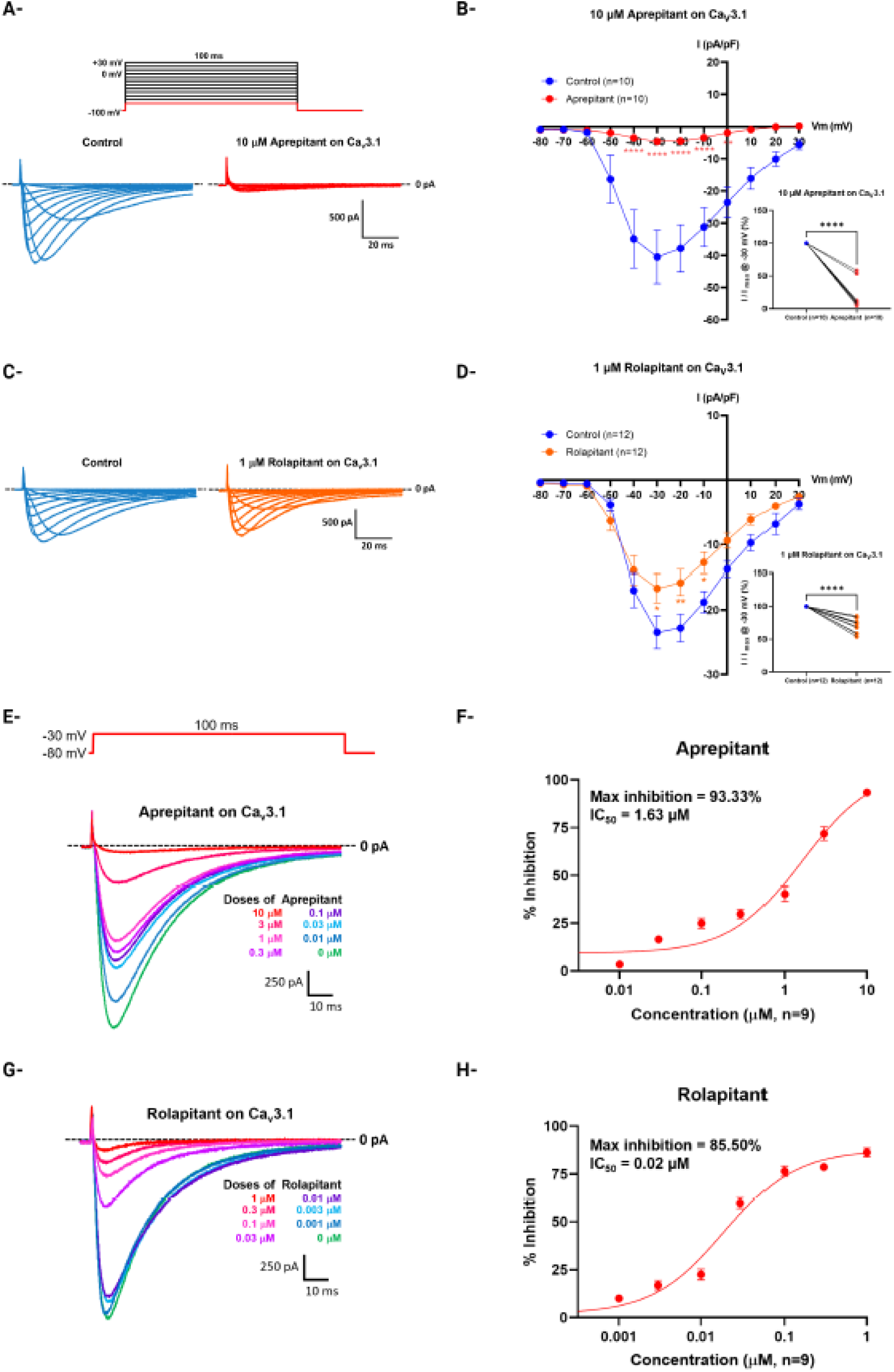
Comparative inhibition of Ca_v_3.1 currents by aprepitant and rolapitant. The selectivity of aprepitant and rolapitant was assessed on the Ca_v_3.1 voltage-gated Ca²⁺ channel transiently expressed in HEK293T cells. Like NALCN, Ca_v_3.1 belongs to the four-domain ion channel superfamily. **A)** and **C)** Representative current traces elicited by a voltage-step protocol in the absence or presence of 10 µM aprepitant and 1 µM rolapitant respectively. **B)** and **D)** Current density–voltage (I–V) relationships for Ca_v_3.1 current derived from **A)** and **C)**. Inset shows paired analysis of current density at −30 mV. The inhibitory properties of both compounds (apparent potency and maximal inhibition) were then characterized using a voltage step to −30 mV from a holding potential of −80 mV. **E)** and **G)** Representative current traces showing Ca_v_3.1 current inhibition following application of increasing concentrations of aprepitant (0.01–10 µM) and rolapitant (0.001–1 µM), respectively. **F)** and **H)** Concentration– response relationships for inhibition of the Ca_v_3.1 current by aprepitant and rolapitant, respectively. Statistical analysis: Two-way ANOVA with Dunnett’s multiple-comparisons test (I–V relationships) and paired t-test (insets panels **B and D**). ****p < 0.0001, ***p < 0.001, **p < 0.01, *p < 0.05.

**Fig. 8:**
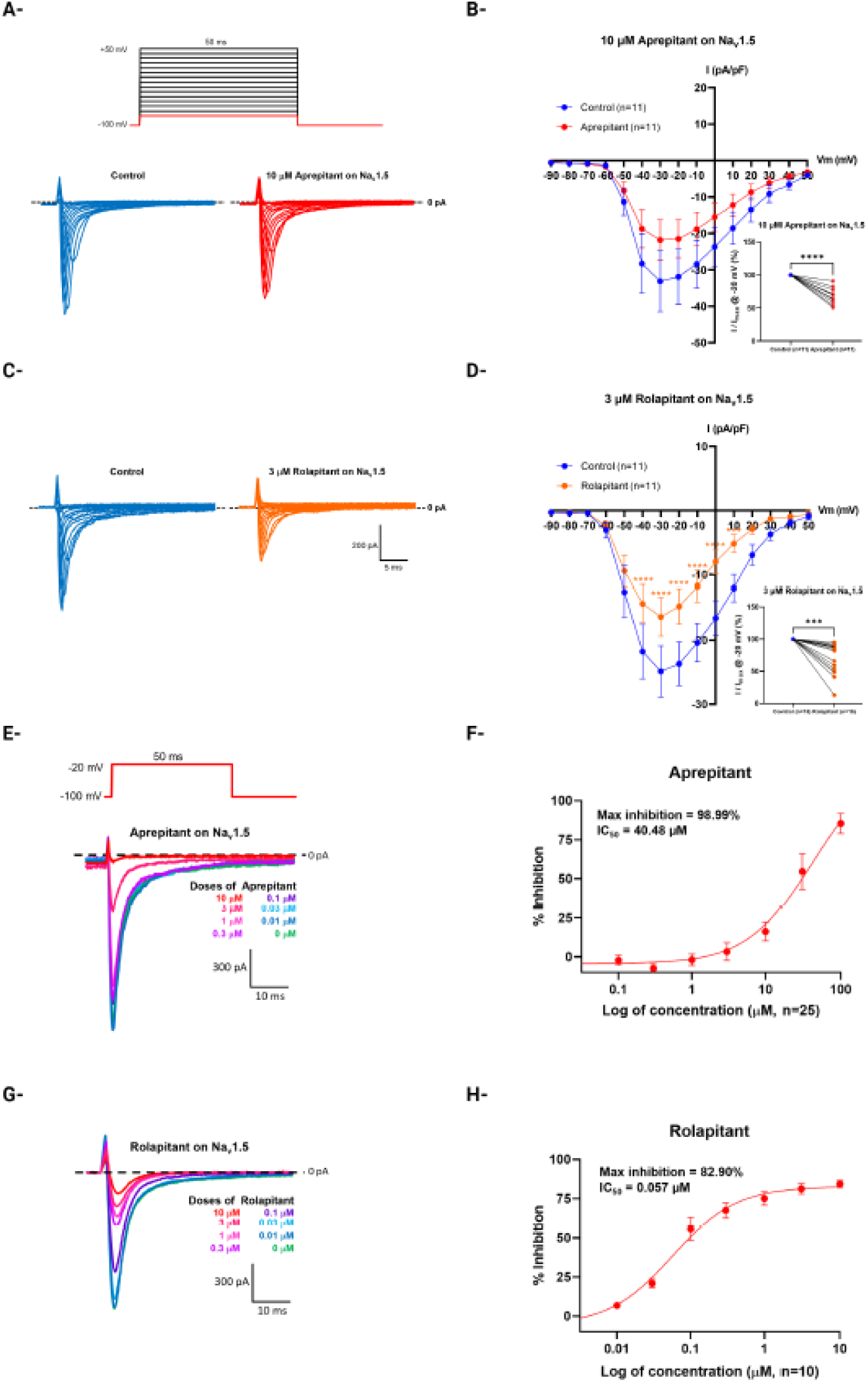
Comparative inhibition of Na_v_1.5 currents by aprepitant and rolapitant. The selectivity of aprepitant and rolapitant was assessed on the Na_v_1.5 voltage-gated Na⁺ channel transiently expressed in HEK293T cells. Like NALCN, Na_v_1.5 belongs to the four-domain ion channel superfamily. **A)** and **C)** Representative current traces elicited by a voltage-step protocol in the absence or presence of 10 µM aprepitant and 3 µM rolapitant respectively. **B)** and **D)** Current density–voltage (I–V) relationships for Na_v_1.5 current derived from **A)** and **C)**. Inset shows paired analysis of current density at −20 mV. The inhibitory properties of both compounds (apparent potency and maximal inhibition) were then characterized using a voltage step to −20 mV from a holding potential of −100 mV. **E)** and **G)** Representative current traces showing Na_v_1.5 current inhibition following application of increasing concentrations (0.01– 100 µM) of aprepitant and rolapitant, respectively. **F)** and **H)** Concentration–response relationships for inhibition of the Na_v_1.5 current by aprepitant and rolapitant, respectively. Statistical analysis: Two-way ANOVA with Dunnett’s multiple-comparisons test (I–V relationships) and paired t-test (insets panels **B and D**). ****p < 0.0001, ***p < 0.001, **p < 0.01, *p < 0.05.

### Electrophysiological Validation in Primary Mouse Hippocampal Neurons

We next examined whether aprepitant modulates the holding current and the resting membrane potential of primary mouse hippocampal neurons, in which both excitability and resting membrane potential have been shown to depend strongly on NALCN[11]. In voltage-clamp recordings at −68 mV, a potential chosen to minimize the contribution of background K⁺ currents, reducing extracellular Ca²⁺ from 2 mM to 0.1 mM produced a substantial increase in holding current, consistent with potentiation of the NALCN current (**Fig. 9A, B**). This current was significantly reduced by 20 µM aprepitant, and was nearly abolished by replacing extracellular Na⁺ with the impermeant cation NMDG⁺. In current-clamp recordings, lowering extracellular Ca²⁺ caused significant membrane depolarization and, in a subset of neurons (5 out of 12), triggered bursting activity (**Fig. 9C-E**). Application of aprepitant (20 µM) markedly hyperpolarized the membrane potential and reduced firing frequency in active cells, an effect that was further enhanced by substitution of extracellular Na⁺ with NMDG⁺ (**Fig. 9E, F**). These observations are consistent with aprepitant inhibiting NALCN-mediated Na⁺ leak currents. Collectively, these results demonstrate that aprepitant inhibits Na⁺ leak currents, resting membrane potential, and neuronal excitability in primary hippocampal neurons. Contributions from additional ion channels or mechanisms cannot be excluded.

**Fig. 9:**
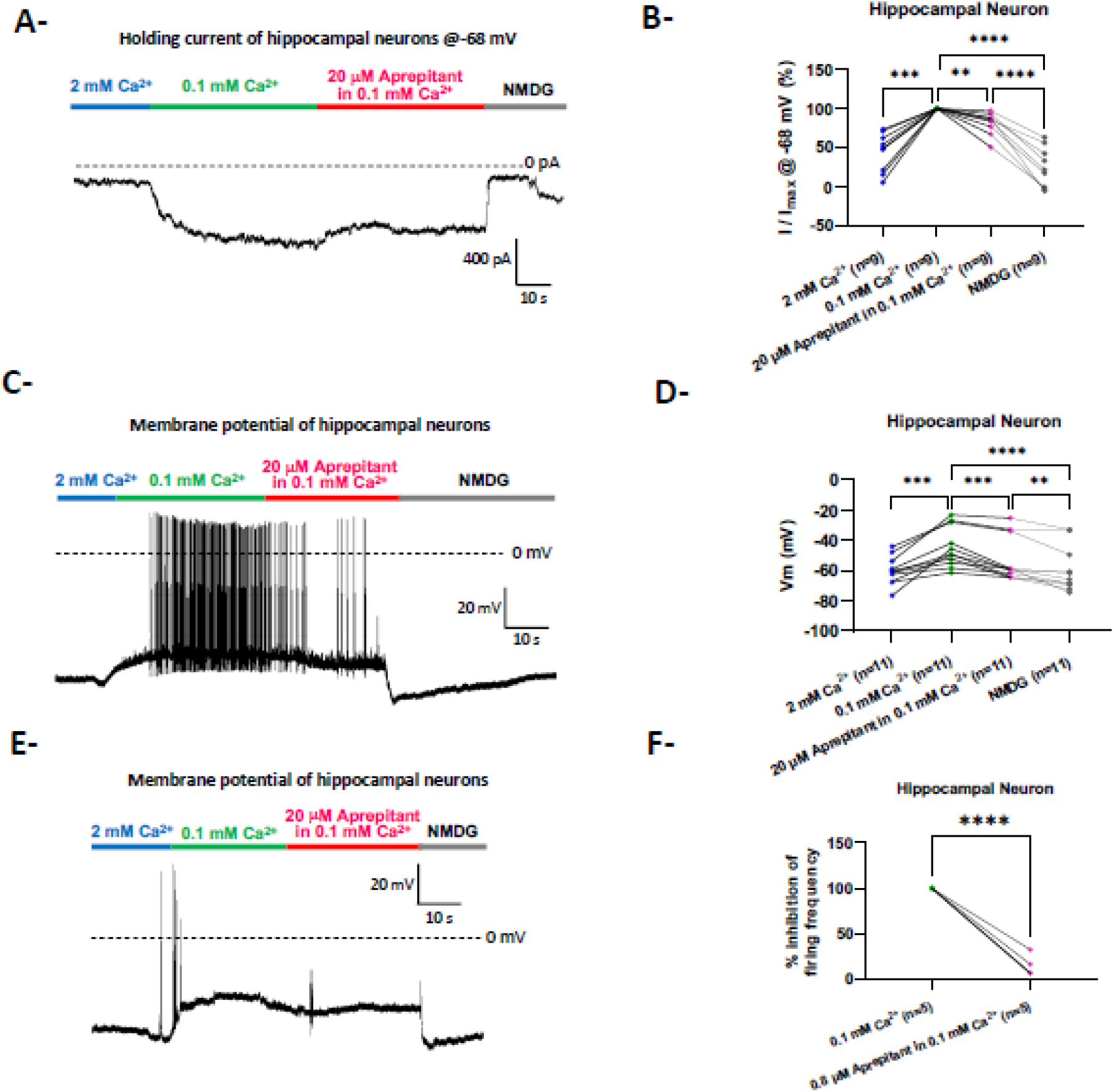
Aprepitant-mediated modulation of electrophysiological properties in mouse primary hippocampal neurons. Whole-cell patch-clamp recordings were performed to assess the effects of aprepitant on both holding current and membrane potential in voltage-clamp and current-clamp configurations, respectively. **A)** Representative voltage-clamp traces from neurons held at −68 mV. Reducing extracellular Ca²⁺ from 2 mM to 0.1 mM produced an increase in holding current, consistent with NALCN current potentiation. This current was partially inhibited by 20 µM aprepitant and was fully abolished upon replacement of extracellular Na⁺ with the large impermeant cation NMDG⁺. **B)** Quantification of changes in holding current under the conditions described in **A)**. **C)** and **E)** Representative current-clamp traces from neurons held at 0 pA. Reducing extracellular Ca²⁺ from 2 mM to 0.1 mM caused membrane depolarization, which in some neurons (*i.e.,* 5 out of 12) was accompanied by spontaneous firing activity**. D)** Membrane potential was hyperpolarized by aprepitant at 20 µM and was further hyperpolarized upon substitution of extracellular Na⁺ with NMDG⁺. **F)** Percentage reduction in firing frequency produced by 20 µM aprepitant under the conditions described in **E)**. Statistical analysis: paired t-test. ****p < 0.0001, ***p < 0.001, **p < 0.01, *p < 0.05.

### In Vivo Validation of Phenotypic Rescue in NALCN Gain-of-Function Model

For *in vivo* validation, we evaluated whether lead repurposing candidates could rescue gain-of-function phenotypes in *C. elegans* carrying a mutation in the NALCN ortholog, *nca-1,* using two established behavioral assays. In the foraging assay, turns and reversals were quantified following removal from bacterial food in the presence of either DMSO (control) or individual test compounds (**Fig. 10A**). Aprepitant and rolapitant significantly reduced the abnormal hyperactive foraging behavior in a dose-dependent manner, while tradipitant had a more modest effect (**Fig. 10A**). In the coiling assay, both aprepitant and rolapitant significantly reduced the frequency of abnormal coiling behavior compared with the DMSO control (**Fig. 10B**). These results demonstrate phenotypic rescue in an *in vivo* NALCN gain-of-function model and are consistent with pharmacological inhibition of NALCN by aprepitant and rolapitant. The differential response among NK-1 receptor antagonists suggests the behavioral effects are unlikely due to a class effect mediated by NK-1 receptor antagonism.

**Fig. 10:**
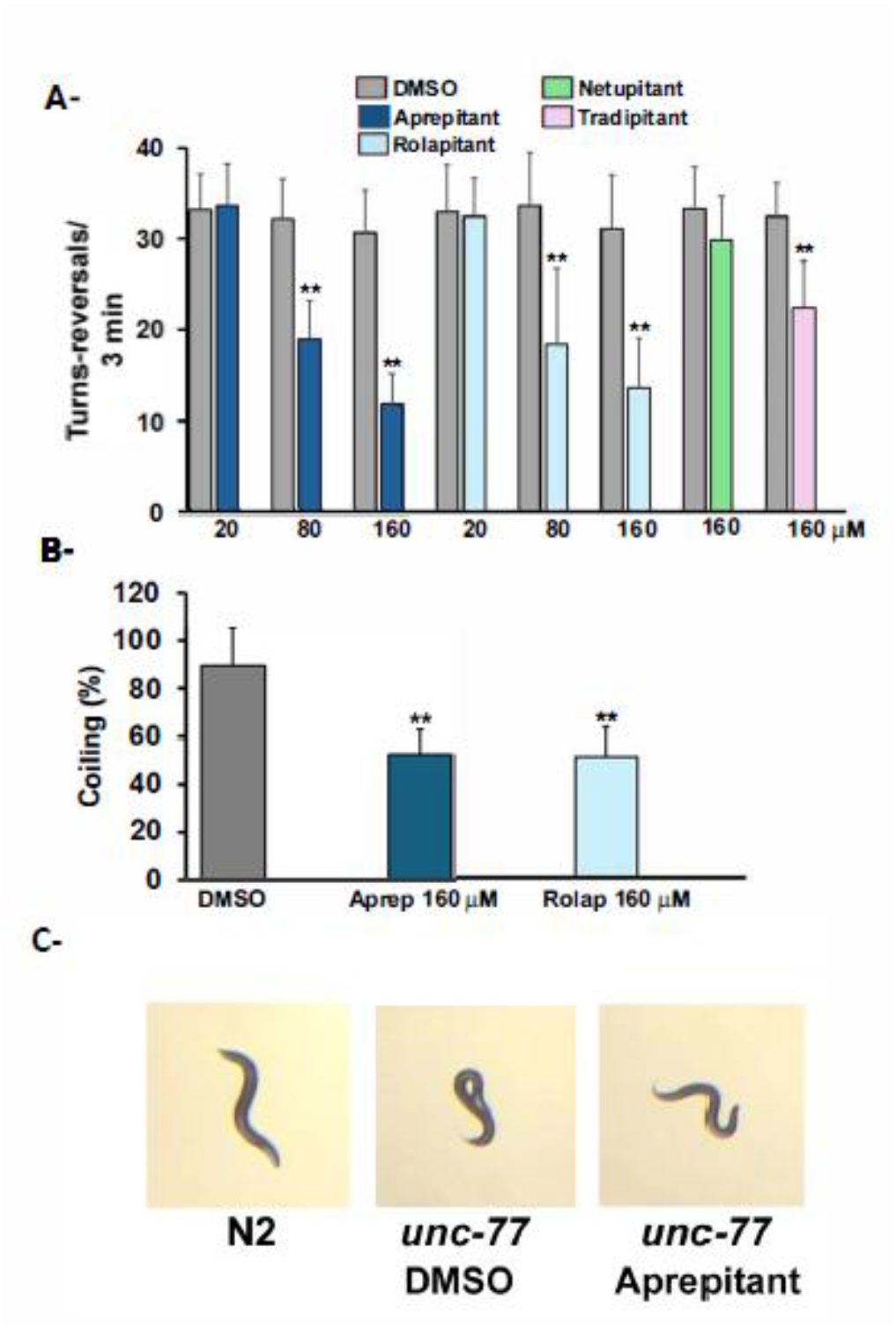
*In vivo* rescue of a NALCN gain-of-function–induced hyperlocomotor phenotype by aprepitant and rolapitant. Aprepitant, rolapitant, netupitant, and tradipitant were tested at the indicated concentrations in *C. elegans unc-77(e625)* mutants. **A)** Locomotor activity expressed as the mean number of movements (turns and reversals) ± SD per condition. Aprepitant, rolapitant, and tradipitant significantly reduced locomotor activity relative to DMSO vehicle control. Netupitant had no significant effect at the concentrations tested. **B)** Percentage of animals displaying coiling behavior, evaluated for aprepitant and rolapitant. Aprepitant and rolapitant significantly reduced coiling compared to DMSO controls. Each experiment was performed at least twice, with a minimum of 10 animals per condition. Vehicle control, low-dose, and high-dose drug treatments were performed on the same day within each experiment. DMSO had no effect on foraging behavior in either wild-type or *unc-77(e625)* animals (data not shown). Statistical analysis: t-test. **p < 0.01. **C)** Representative image of wild-type N2 (left), coiling in DMSO-treated (control) *unc-77* mutants (middle), and reduced coiling in aprepitant-treated *unc-77* mutants.

## Discussion

More than 90% of rare diseases lack approved therapies[5], in large part because conventional drug discovery remains labor-intensive and cost-prohibitive for small patient populations[1,6]. In this study, we established a cost and resource-conscious, AI-enabled translational drug discovery framework integrating phenotypic HTS, structural biology foundation models for more efficient hit-to-lead prioritization, and orthogonal functional validation. Applying this framework for drug repurposing, we identified a lead repurposed candidate for CLIFAHDD, an ultra-rare neurodevelopmental disease with no disease-modifying treatments. AI-based structural modeling reduced a major bottleneck in drug discovery by prioritizing phenotypic screening hits, and there was strong concordance between AI predictions, phenotypic screening, and experimental validation. The findings suggest that NALCN is pharmacologically tractable using small molecules, supporting future therapy development for NALCN channelopathies and more common disorders implicated by NALCN hyperexcitability (*e.g.,* movement disorders such as Parkinson’s disease[54–61]). More broadly, this work demonstrates that integrating empirical and computational approaches can facilitate applying finite experimental resources to the most promising candidates, providing a scalable strategy to accelerate drug repurposing and therapeutic discovery for rare diseases in rapid need of treatments.

### Development and validation of complementary phenotypic high-throughput screening and AI-enabled hit-to-lead prioritization for pharmacological modulators of the NALCN channelosome

While ion channels have historically been successful therapeutic targets[62], the NALCN channelosome has previously been considered a challenging pharmacologic target because functional expression requires a large multiprotein complex and produces a relatively small background current[9]. A prior obstacle has been the lack of a scalable and reproducible recombinant system for phenotypic HTS. We recently addressed this by developing an inducible HEK293 platform to enable reliable, doxycycline-controlled expression of the complete NALCN channelosome[33], Building on this, here we established a fluorescence-based HTS assay capable of quantifying pharmacologic modulation of NALCN activity in 96-and 384-well formats for library-scale screening. Phenotypic hits in this assay were subsequently validated using *in silico, in vitro* and *in vivo* models. This establishes a scalable pipeline for identifying and prioritizing small-molecule modulators of the NALCN channelosome.

Second, NALCN drug discovery has been limited by economic barriers common to rare diseases. A frequent bottleneck in drug discovery is the costly and labor-intensive prioritization of phenotypic screening hits. This bottleneck is particularly impeding for drug repurposing, where bioactive compounds often possess diverse biological functions, and for rare diseases, where drug discovery is increasingly driven by patient advocacy organizations with limited resources[63–66]. Here we employed Boltz-2, a structural biology foundation model, into the discovery pipeline to prioritize compounds based on predicted binding affinity and binding mode. Importantly, we found the AI predictions demonstrated strong concordance with both phenotypic screening and later *in vitro* and *in vivo* experimental validation. Aprepitant was predicted to bind within the NALCN pore through a direct pore-blocking mechanism, which was consistent with electrophysiological findings demonstrating voltage-independent inhibition and sensitivity to extracellular Ca²⁺. Because extracellular Ca²⁺ itself acts as a pore blocker of NALCN[18], the Ca²⁺-dependent shift in apparent potency — with maximal inhibition preserved — supports the compounds occupying an overlapping region of the pore, consistent with the AI-predicted docking model. As noted above, the direction of this effect argues against strict competition for identical sites and instead favors a cooperative relationship, in which Ca²⁺ occupancy stabilizes compound binding. These findings suggest that structural foundation models can streamline hit-to-lead prioritization while generating experimentally testable mechanistic hypotheses.

Beyond hit prioritization, structural modeling also showed promise for hit discovery and lead characterization. Virtual screening of the same FDA-approved library demonstrated substantial concordance with phenotypic HTS, including identification of the eventual lead candidate, aprepitant. These findings suggest that for diseases lacking robust recombinant expression systems or scalable phenotypic assays, AI-based virtual screening may serve as an initial filter to identify hits for targeted experimental validation. Structural modeling further informed structure-activity relationship analyses, correctly predicting aprepitant as the highest-affinity NALCN inhibitor among NK-1 receptor antagonists and anticipating its similar binding profile to NALCN and Ca_v_3.1, but not Na_v_1.5. These predictions were subsequently confirmed by electrophysiology. Collectively, these results support the potential of structural biology foundation models to serve as complementary tools throughout translational drug discovery— from hit discovery and prioritization to mechanistic interpretation and lead optimization.

Overall, these findings suggest that integrating scalable phenotypic assays with structural foundation models can more efficiently identify promising lead candidates, enabling downstream experimental validation to focus on the most promising candidates and minimizing costly testing. Rather than replacing experimental biology, AI served as a force multiplier that amplified the efficiency of empirical discovery and generated testable predictions for later experimental validation. Although demonstrated here for NALCN channelopathy, this framework could be broadly applicable to other diseases with tractable molecular targets and may help lower the time, labor, and financial barriers that have historically limited therapy development for rare diseases.

### NK-1 Receptor Antagonist as Lead Repurposed Drug for NALCN Inhibition

This translational discovery pipeline identified aprepitant as a promising therapeutic candidate for CLIFAHDD, and provided preclinical evidence to support advancement to clinical trials. Phenotypic screening, AI-based structural modeling, electrophysiology, and *C. elegans* results collectively supported aprepitant as a partial NALCN inhibitor and promising therapeutic candidate. Patch-clamp recordings confirmed partial inhibition of NALCN-mediated sodium leak currents (IC₅₀ = 2.22 μM). In primary mouse hippocampal neurons, aprepitant reduced sodium leak current, membrane hyperpolarization, and decreased spontaneous firing frequency. Notably, spontaneous bursting induced by low extracellular Ca²⁺ in a subset of neurons (5 of 12) was decreased by aprepitant. This is consistent with reversal of the neuronal hyperexcitability predicted to result from NALCN gain-of-function variants, which chronically depolarize the resting membrane potential and lower the threshold for spontaneous firing. In a gain-of-function *C. elegans* model, aprepitant produced a dose-dependent rescue of hyperkinetic phenotypes, providing *in vivo* proof-of-concept for pharmacological suppression of NALCN-associated hyperexcitability. These *C. elegans* assays have historically been highly selective among compounds[32], suggesting the behavioral rescue is unlikely to reflect nonspecific locomotor suppression. *C. elegans* lack functional voltage-gated sodium channels and established functional equivalents to the NK-1 receptor or demonstrated sensitivity to substance P [67], suggesting direct modulation of *nca-1*/NALCN and potentially co-expressed T-type calcium channels (highly conserved from nematodes to mammals), as the primary mechanism of behavioral rescue. Together, these results provide convergent evidence supporting aprepitant as a partial inhibitor of NALCN across complementary experimental systems.

Aprepitant is an orally available NK-1 receptor antagonist approved in adults and children age >6 months for chemotherapy-induced nausea and vomiting[68,69]. Prior PET studies demonstrate blood-brain barrier penetration with ≥90% brain NK-1 receptor occupancy at doses ≥100 mg/day or plasma concentrations ≥100 ng/ml, which are achieved at clinically approved doses[40]. This supports the feasibility of achieving brain exposures relevant for partial NALCN inhibition, although the free brain concentration remains due to high plasma protein binding[42,70]. Aprepitant is well-tolerated, and the most common side effects included diarrhea, headache, fatigue, and neutropenia[70,68,69]. While approved for acute use, a prior small study of off-label chronic use in children identified hiccups, fatigue, and headaches as reported side effects[46]. Given the severity of CLIFAHDD and the absence of disease-modifying therapies, the established safety and pharmacokinetic profile of aprepitant make it an attractive candidate for further clinical evaluation. Accordingly, aprepitant has now advanced to a Phase I clinical trial to evaluate the safety and pharmacokinetics of chronic use in individuals with CLIFAHDD (NCT07845825).

Our findings further suggest that NALCN inhibition is not a general property of NK-1 receptor antagonists, but instead depends on specific structural features. Previous studies reported NALCN inhibition by other NK-1 receptor antagonists, including L703606 and maropitant, raising the hypothesis that NALCN is a shared property within this chemical class[33]. In this study, aprepitant and its close structural analog rolapitant demonstrated NALCN inhibition and *in vivo* phenotype rescue, whereas the structurally distinct netupitant and tradipitant showed substantially lower activity. These experimental findings aligned with AI-predicted binding modes, suggesting that the morpholine-acetal scaffold of aprepitant and the spirocyclic lactam core of rolapitant are particularly well suited to occupy the NALCN pore. Together, these results establish an initial structure–activity relationship for NALCN inhibition and provide a rational starting point for future structure-guided medicinal chemistry.

### Pathogenic gain-of-function variants exhibit differing pharmacological sensitivity

Aprepitant exhibited lower maximal inhibition of two CLIFAHDD-associated gain-of-function variants, p.Y578S and p.D600H, than of wild-type NALCN (∼14% versus ∼36% maximal inhibition), extending our previous observation that pathogenic NALCN variants can display altered pharmacologic sensitivity[33]. The mechanisms underlying this differential response remain uncertain. The p.Y578S variant has been shown to increase channel open probability without altering surface expression[26,71], suggesting that altered channel gating could reduce access to the predicted pore-binding site if aprepitant preferentially binds closed or inactivated channel conformations. Alternatively, conformational changes within the pore could reduce drug binding. In contrast, p.D600H retained a similar apparent affinity (IC₅₀ = 2.33 µM) despite reduced maximal inhibition, suggesting that diminished efficacy may primarily reflect altered channel gating rather than impaired binding. These hypotheses warrant future structural and biophysical investigation.

The therapeutic implications of variant-specific pharmacologic sensitivity also merit further study. Because CLIFAHDD typically results from heterozygous gain-of-function variants, patients retain one gain-of-function variant allele and one functional wild-type allele. Selective or preferential modulation of the wild-type channel may therefore reduce the overall pathological sodium leak sufficiently to restore physiological neuronal excitability without requiring more potent channel blockage of the mutant variant. Whether this potential “heterozygous rescue” translates into clinical benefit will be evaluated in future clinical studies.

### Selectivity profiling reveals possible dual therapeutic opportunity

Selectivity profiling of aprepitant and rolapitant against two additional members of the four-transmembrane domain ion channels revealed distinct inhibitory profiles. Aprepitant exhibited relatively weak inhibition of Na_v_1.5 (IC₅₀ ≈ 40 μM), approximately 18-fold less potent than NALCN. Rolapitant inhibited Na_v_1.5 with substantially higher potency (IC₅₀ ≈ 0.057 μM), representing a potential selectivity liability that may increase the risk of cardiac adverse effects.

Alternatively, Ca_v_3.1 inhibition may represent a therapeutic opportunity rather than simply an off-target effect. Aprepitant inhibited Ca_v_3.1 with potency similar to NALCN, while rolapitant was markedly more potent against Cav3.1 (approximately 172-fold higher), suggesting this channel may be its primary ion channel target. Similar to NALCN, gain-of-function variants in CACNA1G (encoding Ca_v_3.1) are associated with cerebellar ataxia and progressive atrophy, including spinocerebellar ataxia type 42 and an ultrarare developmental and epileptic encephalopathy[72–75]. Thalamic expression of Ca_v_3.1 suggest it as a pharmacological target for absence epilepsy[76]. Dual inhibition of NALCN and Ca_v_3.1 could provide convergent suppression of two distinct depolarizing mechanisms, the background sodium leak and the low-voltage-activated calcium influx, potentially producing an additive or synergistic anti-hyperexcitability effect, though this has not yet been tested directly. Whether this dual pharmacology confers additional benefit for CLIFAHDD or other disorders of neuronal hyperexcitability warrants further evaluation.

### Limitations and future opportunities

This study has several limitations. First, in this study we prioritized compounds predicted to bind to the NALCN central pore to inhibit NALCN *via* direct pore blockade. Hits from phenotypic screening may act as NALCN antagonists *via* other mechanisms, and should be evaluated in future studies. Second, electrophysiology characterization of aprepitant and rolapitant on Ca_v_3.1 and Na_v_1.5 was conducted at single membrane potentials, limiting evaluation of whether inhibition is tonic, use-dependent, or voltage-dependent. This can be assessed in follow-up studies using multi-pulse voltage protocols. Third, pharmacological characterization of NALCN gain-of-function variants was limited to two CLIFAHDD-associated variants (p.Y578S and p.D600H) and may not generalize to other *de novo* variants. Fourth, ion channel profiling was restricted to Ca_v_3.1 and Na_v_1.5, and broader profiling would strengthen the selectivity characterization. Fifth, while the *C. elegans in vivo* data provide proof-of-concept for pharmacological rescue of NALCN gain-of-function hyperexcitability, the internal drug concentrations achieved *in vivo* remain uncertain. This should be evaluated in future mammalian models, such as knock-in mice expressing pathogenic gain-of-function NALCN variants (currently in development). Sixth, the primary screen was a single-pass, single-concentration (10 µM) phenotypic screen using a 60% suppression threshold, and further assay optimization is necessary for broader drug screening. Seventh, concentration-response relationships were determined only for human NALCN expressed in HEK293 cells. The primary hippocampal neuron experiments used a single concentration of aprepitant (20 µM), and the *C. elegans* experiments used compound applied to agar plates (20-160 µM), where internal exposure is unknown. While the mouse and nematode results support target engagement and phenotypic consequence, the potency across species remains unclear.

Looking forward, the AI-enabled discovery framework described here can be applied to larger and more chemically diverse compound libraries. Coupling AI-guided virtual screening with scalable phenotypic HTS provides a practical strategy for identifying next-generation NALCN modulators with improved potency, selectivity, and drug-like properties. Structure-guided medicinal chemistry based on the predicted binding mode of aprepitant may further optimize NALCN selectivity or, alternatively, utilize dual NALCN/Ca_v_3.1 inhibition. Finally, patient-derived hiPSC neurons and genetically engineered animal models in development will be valuable for validating efficacy, defining pharmacodynamic biomarkers, and prioritizing compounds for clinical translation.

## Conclusions

In summary, we established a cost and resource-conscious, AI-enabled translational drug discovery framework that integrates scalable phenotypic screening, structural biology foundation models, and orthogonal functional validation to accelerate therapeutic discovery for rare diseases. Applying this framework to CLIFAHDD, an ultra-rare channelopathy with no approved treatments, we identified aprepitant as a promising therapeutic candidate for future clinical translation, and demonstrated NALCN as a pharmacologically tractable target for small molecules. The study provides a roadmap for integrating computational models into translational drug discovery to reduce costs and improve efficiency in hit-to-lead prioritization with demonstrated experimental validation. More broadly, these findings support AI-assisted drug repurposing as a scalable strategy to reduce common economic and biologic barriers in drug repurposing and therapy discovery, ultimately making therapies more attainable for the growing number of people affected by rare and ultra-rare diseases.

## Declarations

### Ethics approval and consent to participate

All animal procedures were conducted in accordance with the European Communities Council Directive and were reviewed and approved by the IGF institute’s local Animal Welfare Unit (A34-172-41).

### Consent for publication

Not Applicable

### Availability of data and materials

The datasets used and/or analyzed during the current study are available from the corresponding authors on reasonable request. The cell lines and other research materials generated in this study are available from corresponding authors upon reasonable request.

### Authors’ contributions

NT contributed to study conceptualization, patch-clamp data curation, methods, and analyses, visualization; and manuscript writing and editing. JAT contributed to conceptualization, method and drug development framework, interpretation, supervision, funding acquisition, and manuscript writing and editing. SP contributed to high-throughput screening assay development and optimization, data acquisition, interpretation, visualization, and manuscript editing. AT contributed to patch-clamp investigations, analysis, and manuscript editing. NC contributed to assay development and results interpretation. BH contributed to HTS assay development, study design, methods, and manuscript review. PL contributed to manuscript review and editing. DSD contributed to study design and methodology, *C. elegans* model data acquisition, analyses, interpretation, and visualization, resources, and manuscript writing and editing. DZ contributed to study methodology, analyses, interpretation, resources, supervision, and manuscript editing. AM contributed to study conceptualization, methodology, data curation, analyses, critical interpretation, funding acquisition, resources, supervision, validation, and manuscript writing and editing. YY contributed to conceptualization, method development, high-throughput screening and AI-based modeling data curation, analyses, interpretation, and visualization, funding acquisition, and manuscript writing and editing. All authors read and approved the final manuscript.

## Acknowledgements

We express our gratitude to the Channeling Hope Foundation (https://www.channelinghope.org/), a 501c3 non-profit patient advocacy group for NALCN Channel Related Diseases, that kindly provided financial support to NT and facilitated the drug discovery collaboration. We thank Dr. Jacinthe Gingras for serving as a scientific advisor for drug repurposing. We thank the technical support from the Nathan Shock Center San Antonio funded by P30AG013319 and the Target Discovery Core and Center for Innovative Drug Discovery funded by grants from National Cancer Institute (P30CA54174) and Cancer Prevention and Research Institute of Texas (CPRIT) grants (RP210208 and RP250601). The authors acknowledge the Texas Advanced Computing Center (TACC) at The University of Texas at Austin for providing computational resources that have contributed to the research results reported within this paper (http://www.tacc.utexas.edu).

## Abbreviations

AI: Artificial Intelligence
Cav: Voltage-gated Ca2+ channel
CLIFAHDD: Congenital contractures of the limbs and face, hypotonia, and developmental delay
CNS: Central nervous system
DMSO: Dimethyl sulfoxide
Dox: Doxycycline
FAM155A/B: Family with Sequence Similarity 155 Member A/B
FDA: Food and Drug Administration
FLIPR: Fluorescence Imaging Plate Reader
HEK293: Human Embryonic Kidney 293
HTS: High-throughput screening
NALCN: Na+ leak channel
Nav: Voltage-gated Na+ channel
NK-1: Neurokinin-1
STR: Short Tandem Profiling
UNC79: Uncoordinated-79 Homolog
UNC80: Uncoordinated-80 Homolog

