## Supplementary material for "An AI-Enabled Translational Drug Discovery Framework for NALCN Channelopathy: From High-Throughput Screening to Therapeutic Candidate": Supplemnentary Figures

### Supplementary Figures

#### Increasing depolarization with increasing doxycycline

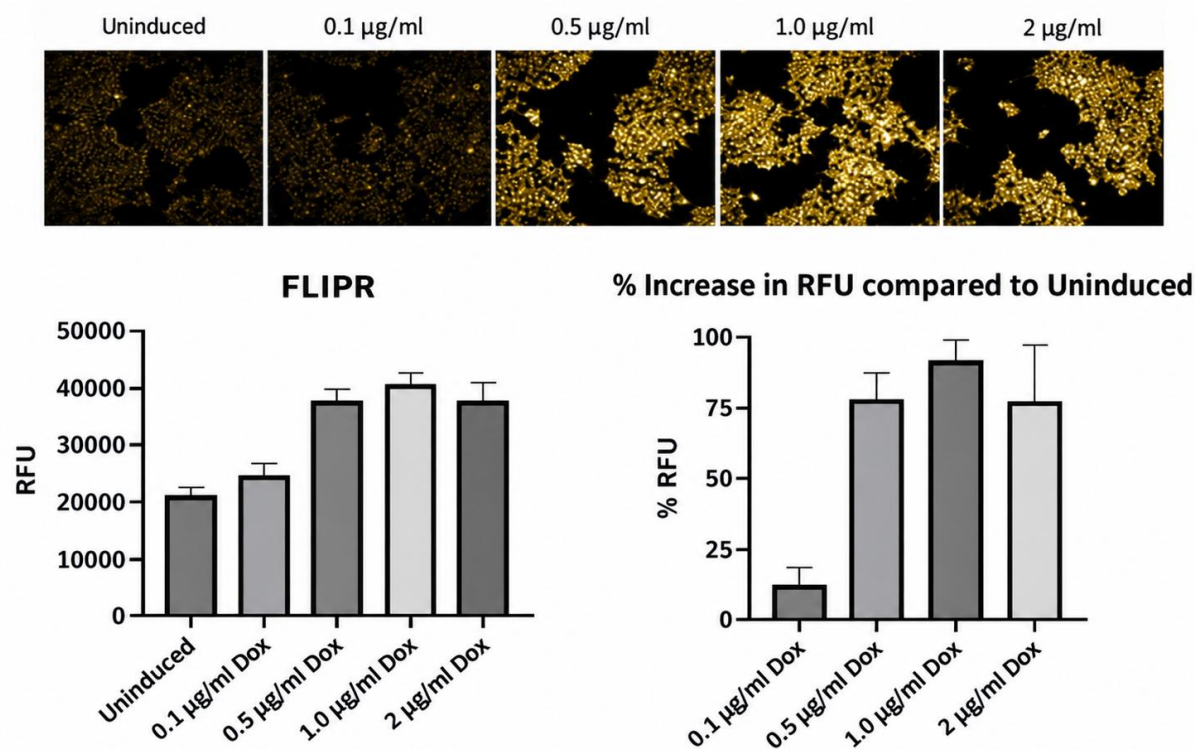

**Fig. S1: Effects of various Dox concentrations on membrane depolarization in HTS assay.** The optimized concentration was determined to be 1.0 µg/mL.

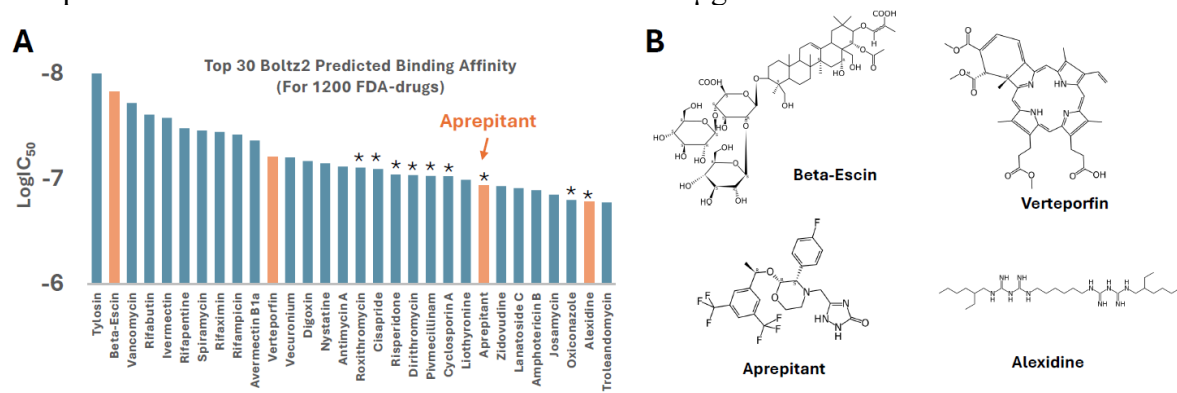

**Fig. S2: Boltz2-predicted binding affinities of 1,200 FDA-approved drugs to NALCN. A)** The top 30 predicted candidates. The top 4 hits identified from the phenotypic high-throughput screening (HTS) are highlighted in orange. An asterisk (\*) denotes molecules predicted to bind within the central pore of NALCN. **B)** Chemical structures of the top 4 HTS hits.

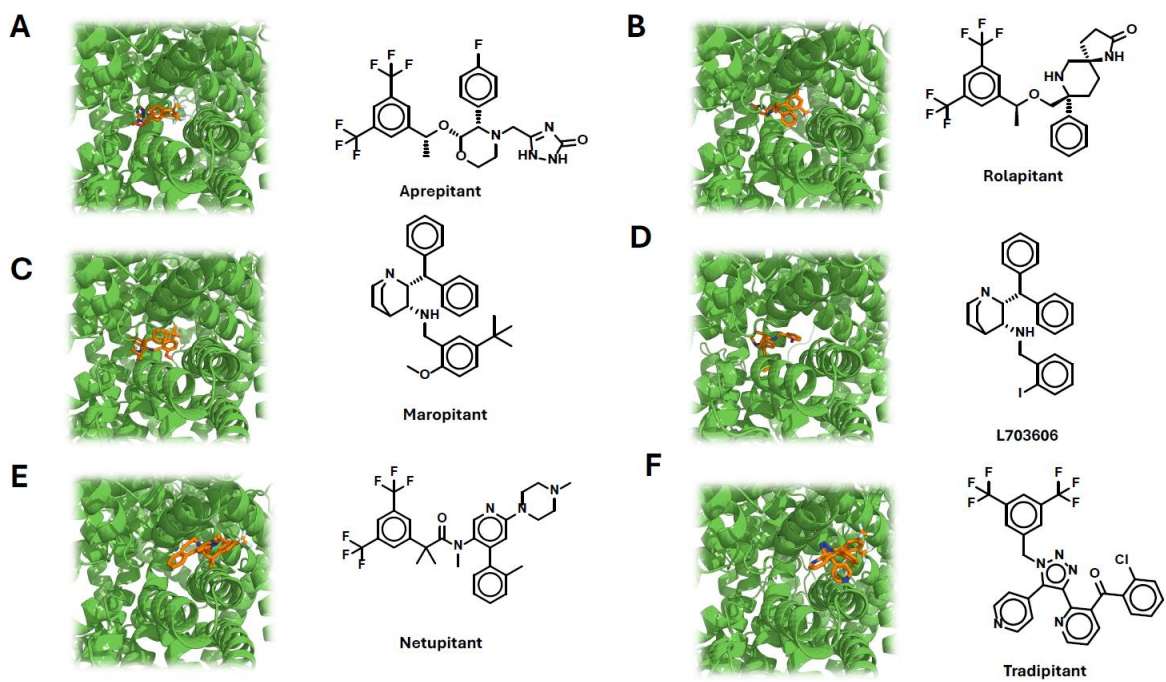

**Fig. S3: Boltz-2 predicted binding modes of NK-1 Receptor Antagonists.** A-D) Aprepitant, Rolapitant, Maropitant and L703606 bind to the central pore of NALCN. E, F) Netupitant and Tradipitant bind to the entry region of NALCN

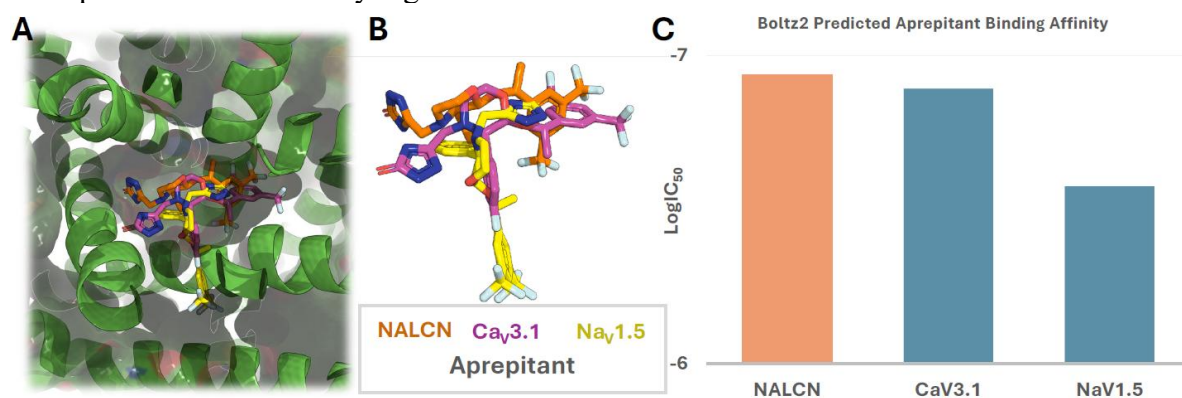

**Fig. S4: Analysis of the binding modes of Aprepitant with CaV3.1 and NaV1.5.** A) Superimposition of the binding modes of Aprepitant within NALCN, CaV3.1, and NaV1.5. B) Ligand-only view of the superimposed conformations. C) Binding energies predicted by Boltz2.

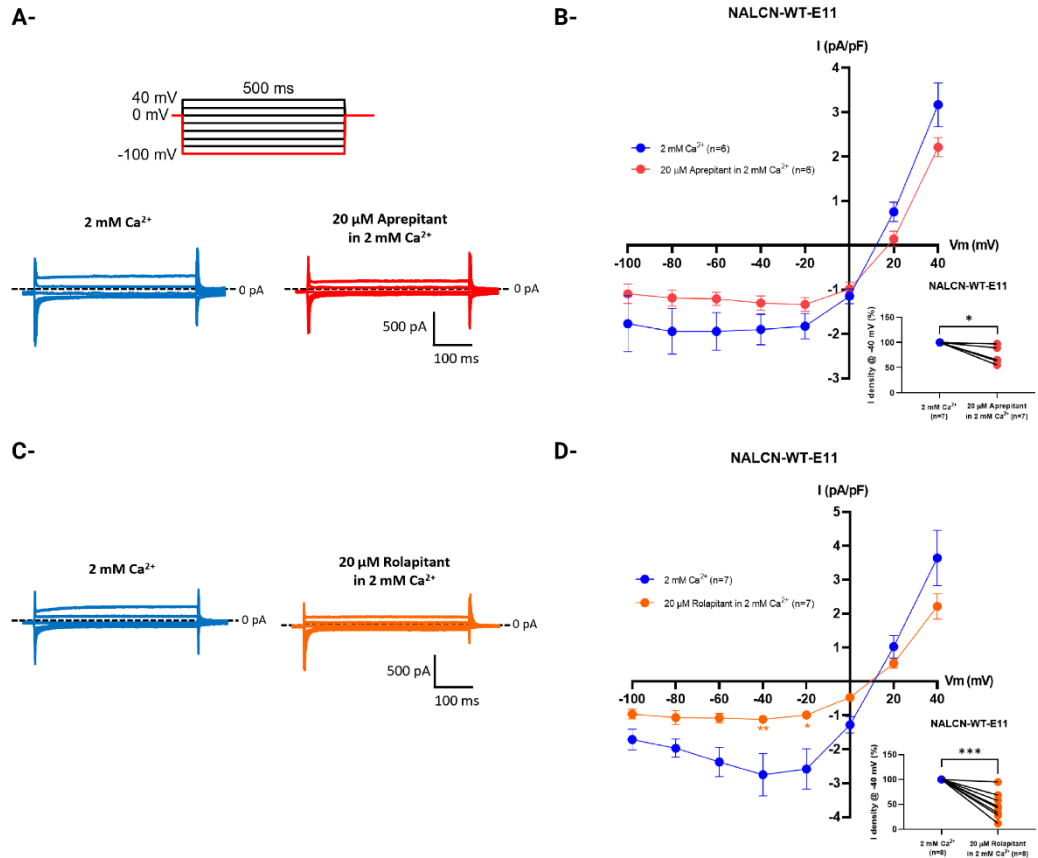

**Fig. S5: Modulation of the NALCN-mediated $\text{Na}^+$ leak current by aprepitant and rolapitant in 2mM extracellular $\text{Ca}^{2+}$.** The effects of aprepitant and rolapitant were assessed at 20 $\mu\text{M}$ on the NALCN current elicited by a voltage-step protocol in the monoclonal HEK293 cell line stably expressing the wild-type NALCN channelosome [33]. **A)** and **C)** Representative current traces recorded in 2 mM extracellular $\text{Ca}^{2+}$ before and after application of aprepitant and rolapitant, respectively. **B)** and **D)** Current density–voltage (I/V) relationships derived from **A)** and **C)**, respectively. Insets show paired analyses of current density at $-40$ mV. Statistical analysis: two-way ANOVA with Dunnett's multiple comparisons test (I/V relationships) and paired t-test (insets). \*\*\*\*$p < 0.0001$, \*\*\*$p < 0.001$, \*\*$p < 0.01$, \*$p < 0.05$.

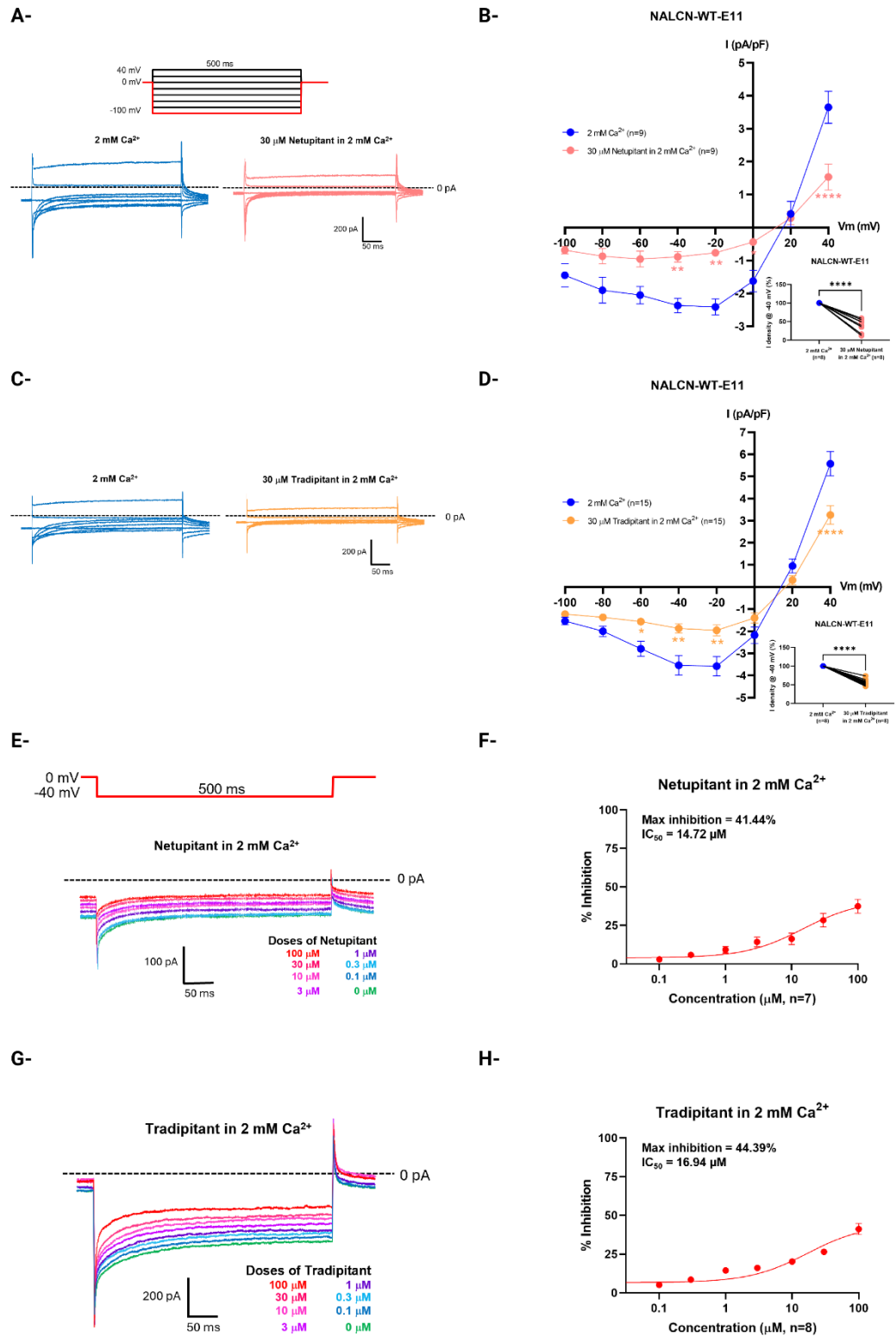

**Fig. S6: Concentration–response characterization of NALCN current inhibition by netupitant and tradipitant in 2mM extracellular $\text{Ca}^{2+}$.** The effects of netupitant and tradipitant were first assessed at 20 $\mu\text{M}$ on the NALCN current elicited by a voltage-step protocol in the monoclonal HEK293 cell line stably expressing the wild-type NALCN channelosome [33]. Then the inhibitory properties of netupitant and tradipitant on the NALCN current (apparent affinity and maximal inhibition) were characterized using a voltage step to

–40 mV from a holding potential of 0 mV. **A)** and **C)** Representative current traces recorded in 2 mM extracellular $\text{Ca}^{2+}$ before and after application of netupitant and tradipitant, respectively. **B)** and **D)** Current density–voltage (I/V) relationships derived from **A)** and **C)**, respectively. Insets show paired analyses of current density at –40 mV. **E)** and **G)** Representative current traces showing NALCN current inhibition following application of increasing concentrations (0.1–100 $\mu\text{M}$) of netupitant and tradiptant, respectively. **F)** and **H)** Concentration–response relationships for inhibition of the NALCN current by netupitant and tradiptant, respectively. Statistical analysis: two-way ANOVA with Dunnett's multiple comparisons test (I/V relationships) and paired t-test (insets). \*\*\*\*$p < 0.0001$, \*\*\*$p < 0.001$, \*\*$p < 0.01$, \*$p < 0.05$.

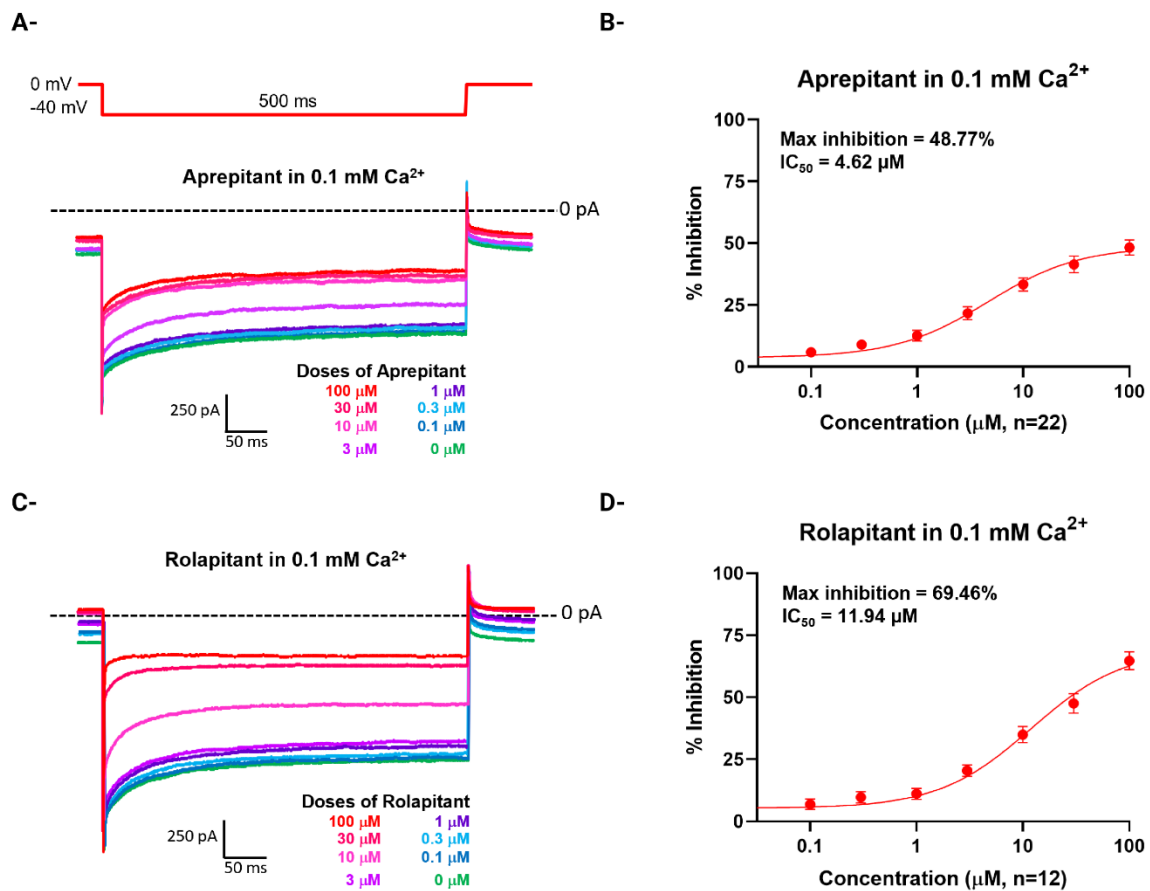

**Fig. S7: Concentration–response characterization of NALCN current inhibition by aprepitant and rolapitant in 0.1mM extracellular $\text{Ca}^{2+}$.** The inhibitory properties of aprepitant and rolapitant on the NALCN current (apparent affinity and maximal inhibition) were characterized using a voltage step to –40 mV from a holding potential of 0 mV. **A)** and **C)** Representative current traces showing NALCN current inhibition following application of increasing concentrations (0.1–100 $\mu\text{M}$) of aprepitant and rolapitant, respectively. **B)** and **D)** Concentration–response relationships for inhibition of the NALCN current by aprepitant, and rolapitant, respectively.
