## Supplementary Materials for "An AI-Enabled Translational Drug Discovery Framework for NALCN Channelopathy: From High-Throughput Screening to Therapeutic Candidate"

### Supplementary Methods

#### *In Vitro Validation with patch clamp recordings*

The lead candidate and mechanistically related compounds (NK-1 receptor antagonists) were advanced to orthogonal functional validation by whole-cell patch clamp electrophysiology. In vitro validation was designed to assess (a) NALCN current inhibition in HEK293 cells stably expressing the wild-type NALCN channelosome; (b) generalizability by evaluating NALCN current inhibition in HEK293 cells stably expressing CLIFAHDD patient-derived gain-of-function variants; (c) selectivity by evaluating compound effects on Ca<sub>v</sub>3.1 and Na<sub>v</sub>1.5 currents in HEK293 cells; (d) inhibition of endogenous NALCN current in mouse primary hippocampal neurons.

#### *Plasmids*

The expression plasmid to express the Ca<sub>v</sub>3.1 T-type voltage-gated calcium channel was previously described (Monteil et al, 2000). The expression plasmid to express the Na<sub>v</sub>1.5 voltage-gated sodium channel was kindly provided by Dr Céline Marionneau. The pcDNA3-GFP plasmid was previously described (Swayne et al, 2009). Integrity of each plasmid was carefully checked by DNA sequencing following amplification (Eurofins Genomics).

#### *Transfection*

For functional expression of Nav1.5 and Cav3.1 channels, 1.9 µg of plasmid DNA supplemented with 0.1 µg of pcDNA3-GFP was used to transfect HEK293T cells in 35 mm Petri dishes using the jetPEI reagent (Polyplus #101000053) according to the manufacturer's protocol.

#### *Chemical compounds*

Compounds identified and assayed in this study were Rolapitant (Selleckchem #Z815946-5 mg), Aprepitant (Sigma-Aldrich #SML2215-5 mg), Netupitant (MedChemExpress # HY-16346) and Tradipitant (MedChemExpress #HY-16732). All other chemicals were purchased from Sigma-Aldrich.

#### *Patch clamp recordings of HEK293 and HEK293T cells*

Cells were dissociated with Versene (Thermo Fisher Scientific #15040066). Macroscopic sodium leak (NALCN), Cav3.1 and Nav1.5 currents were recorded at room temperature in the whole cell configuration using an Axopatch 200B amplifier (Molecular Devices). Data were acquired using the pCLAMP10 software (Molecular Devices) and an Axon Digidata 1440A digitizer (Molecular Devices) at 10 kHz. Patch pipettes were pulled from 1.5/086 borosilicate glass capillaries (Sutter Instrument CO. #BF150-86-15). For NALCN currents, intracellular solution contained 140 mM CsCl, 10 mM CsF, 5 mM EGTA, 10 mM HEPES and 2 mM Na<sub>2</sub>ATP, pH was adjusted at 7.2 with CsOH, ~ 285 mosm/L. Extracellular solution contained 150 mM NaCl, 2 mM CaCl<sub>2</sub>, 1.2 MgCl<sub>2</sub>, 10 mM HEPES, 10 mM D-(+)-glucose, pH 7.4 adjusted with NaOH, ~ 305 mosm/L. Other extracellular solutions used were (1) NMDG 150 mM, CaCl<sub>2</sub> 2 mM, MgCl<sub>2</sub> 1.2 mM, HEPES 10

mM, D-(+)-glucose 10 mM, the pH was adjusted at 7.4 with HCl, ~ 315 mosm/L (2) NaCl 150 mM, CaCl<sub>2</sub> 0.1 mM, MgCl<sub>2</sub> 1.2 mM, HEPES 10 mM and D-(+)-glucose 10mM, pH 7.4 adjusted with NaOH, ~ 300 mosm/L.

For Ca<sub>v</sub>3.1 currents, intracellular solution contained 140 mM CsCl, 10 mM EGTA, 10 mM HEPES, 3 mM CaCl<sub>2</sub>, 0.6 mM NaGTP and 3 mM MgATP. pH was adjusted at 7.2 with CsOH, ~ 285 mosm/L. Extracellular solution contained 135 mM NaCl, 20 mM TEACl, 2 mM CaCl<sub>2</sub>, 1 mM MgCl<sub>2</sub>, 10 mM HEPES, pH 7.4 adjusted with NaOH, ~ 305 mosm/L.

For Na<sub>v</sub>1.5 currents, intracellular solution contained 5 mM KCl, 125 mM CsCl, 5 mM TEACl, 2 mM NaCl, 0.5 mM EGTA, 10 mM HEPES, 3 mM MgATP and 0.3 mM NaATP. pH was adjusted at 7.2 with CsOH, ~ 285 mosm/L. Extracellular solution contained 140 mM NaCl, 5.6 mM KCl, 2 mM CaCl<sub>2</sub>, 0.8 mM MgCl<sub>2</sub>, 10 mM HEPES and 5 mM D-(+)-glucose, pH 7.4 adjusted with NaOH, ~ 305 mosm/L.

#### *Cultures of mouse primary hippocampal neurons*

Primary hippocampal neurons were prepared from newborn C57BL/6J pups as previously described (Moutin et al, 2020). Hippocampi were dissected under a stereomicroscope and transferred into Hibernate-A medium (Thermo Fisher Scientific #A1247501) supplemented with 1% penicillin/streptomycin (Thermo Fisher Scientific #15140122). Dissected tissue was enzymatically digested with papain (50 U/mL; Sigma-Aldrich #76616) for 10 min at 37 °C, followed by a 5-min co-incubation with DNase I (Roche #11284932001). Enzymatic digestion was terminated by the addition of complete growth medium composed of Neurobasal-A (86.5%; Thermo Fisher Scientific #10888022) supplemented with fetal bovine serum (10%; Thermo Fisher

Scientific #10270106), penicillin/streptomycin (1%; Thermo Fisher Scientific #15140122), B27™ Plus supplement (2%; Thermo Fisher Scientific, #A3582801), L-glutamine (0.25%; Thermo Fisher Scientific #25030024), and GlutaMAX™ (0.25%; Thermo Fisher Scientific #35050061).

Mechanical dissociation was achieved by gentle trituration in three successive rounds of decreasing intensity, collecting the supernatant after each round. The pooled supernatant was carefully layered onto a bubble cushion (4% bovine serum albumin in Hibernate-A, Sigma-Aldrich #A7906, supplemented with 10% penicillin/streptomycin) and centrifuged at $300 \times g$ for 7 min. The resulting cell pellet was resuspended in complete growth medium and plated onto 35-mm culture dishes that had been sequentially coated with poly-L-ornithine (0.03 mg/mL in PBS, overnight) and laminin (Sigma-Aldrich #L2020), 1 µg per dish in Neurobasal-A for at least 2 h).

Cells were plated at a density of 150,000 cells per dish in 2 mL of complete growth medium (two dishes per pup) and maintained at 37 °C in a humidified incubator with 5% CO<sub>2</sub>. On day *in vitro* 2 (DIV2), cytosine-β-D-arabinofuranoside (1 µM ; Sigma-Aldrich #C6645) was added to inhibit glial proliferation. On DIV3, 80% of the medium was replaced with serum-free BrainPhys™ medium (STEMCELL Technologies #05790) supplemented with B27™ Plus supplement (2%; Thermo Fisher Scientific #A3582801), GlutaMAX™ (0.25%; Thermo Fisher Scientific #35050061), and penicillin–streptomycin (1%; Thermo Fisher Scientific #15140122). On DIV6, a partial medium change was performed to support continued neuronal maturation.

*Patch clamp recordings of mouse primary hippocampal neuron*

Hippocampal cultures from >3 independent litters were used for electrophysiological recordings between DIV14 and DIV20, after synapse maturation. Intracellular solution contained 10 mM NaCl, 12 mM KCl, 118 mM K-gluconate, 10 mM HEPES, 4 mM MgCl<sub>2</sub>, 5 mM Tris, 0.3 mM Tris-GTP, 14 mM, Tris-creatine phosphate, and 4 mM MgATP. Extracellular solution contained 140 mM NaCl, 10 mM HEPES, 10 mM D-glucose, 3 mM KCl, 2 mM MgCl<sub>2</sub>, and 2 mM CaCl<sub>2</sub>. Other extracellular solutions that were used contained (1) 140 mM NMDG, 10 mM HEPES, 10 mM D-glucose, 3 mM KCl, 2 mM MgCl<sub>2</sub>, and 2 mM CaCl<sub>2</sub>. and (2) 140 mM NaCl, 10 mM HEPES, 10 mM D-glucose, 3 mM KCl, 2 mM MgCl<sub>2</sub>, and 0.1 mM CaCl<sub>2</sub>. pH was adjusted at 7.4 with NaOH, ~ 300 mosm/L for all solutions.

##### *In Vivo Validation in C. elegans with gain-of-function mutation in NALCN*

For *in vivo* orthogonal validation, we used a *C. elegans* unc-77(e625) model, which harbors a gain-of-function mutation in the NALCN ortholog, *nca-1*[32]. These variants display a reproducible gain-of-function phenotype characterized by hyperactive locomotion, exaggerated body bending, and excessive coiling. This phenotype has been previously validated as a quantitative pharmacodynamic readout of NALCN inhibition[32].

##### *Drug Concentrations for C. elegans*

Drug concentrations were selected based on prior *C. elegans* studies of NK-1 receptor antagonists by our group [33] and psychotropic drugs [37,38]. The affinity of various drugs for *C. elegans* receptors are similar to their human counterparts (in the high nanomolar range) measured by direct binding to receptors expressed in cell lines. However, micromolar levels (20-160 μM) of drug are required on agar plates to produce physiological responses mediated by these same receptors. The roughly hundred-fold difference between *in vitro* (plate) and *in vivo* concentrations is attributed to limited uptake through the cuticle and restricted tissue distribution [39].

*Foraging Assay:* We used a previously described assay to measure the hyperactive foraging response of *unc-77* mutants with NALCN gain-of-function to the lead candidate and other NK-1 receptor antagonists [32]. Briefly, we prepared small (60 mm diameter) seeded agar plates by coating them with control buffer (dilute acetic acid [$10^{-4}$ M] with dimethyl sulfoxide, DMSO, to match the drug conditions) or drug dissolved in DMSO and then diluted in acetic acid buffer. Animals were transferred to control or drug plates for 2 hours to allow uptake and equilibration of drugs. To evaluate the foraging response, animals were transferred from the original plates with food to large agar plates that lacked bacteria. We then observed foraging behavior and counted the total number of sharp “omega” turns (head touching body), pirouettes, and reversals made over 3 min. Each experiment was repeated at least twice, and data was obtained from a minimum of 10 animals.
